# lncRNAs contribute to caste differentiation as a regulatory layer in ants

**DOI:** 10.64898/2026.01.08.698392

**Authors:** Guo Ding, Fuqiang Lin, Jixuan Zheng, Dashuang Zuo, Zijun Xiong, Chenyan Liao, Bitao Qiu, Wenjiang Zhong, Jie Zhao, Weiwei Liu, Guojie Zhang

**Affiliations:** Center for Evolutionary & Organismal Biology, Women’s Hospital & Liangzhu Laboratory, Zhejiang University School of Medicine, Hangzhou, 310058, China; State Key Laboratory of Genetic Evolution & Animal Models, Kunming Institute of Zoology, Chinese Academy of Sciences, Kunming, 650201, China; BGI Research, Wuhan, 430074, China; Section for Ecology and Evolution, Department of Biology, University of Copenhagen, Copenhagen, 2100, Denmark; Evolutionary Biology & Ecology, University of Freibury, 79104, Germany; Yunnan Key Laboratory of Biodiversity Information, Kunming Institute of Zoology, Chinese Academy of Sciences, Kunming, 650201, China

## Abstract

Caste differentiation in ants represents one of the major evolutionary transitions and provides a unique model for studying developmental mechanisms underlying division of labor. Genetically identical individuals follow divergent epigenetically-regulated developmental trajectories that give rise to morphologically distinct phenotypes and specialized roles. While long non-coding RNAs (lncRNAs) are emerging as key epigenetic regulators across diverse biological systems, their specific contributions to caste development in social insects remain largely unexplored. Here, we conducted comprehensive transcriptomic analyses across major developmental stages of two evolutionarily and ecologically distinct ants Monomorium pharaonis and Acromyrmex echinatior, identifying over 10,000 lncRNAs. We demonstrated that lncRNAs exhibit dynamic, caste-specific expression patterns throughout development. We also identified a subset of lncRNAs displaying canalized expression patterns, characterized by progressively increasing caste bias and decreasing within-caste variation as development proceeds. Co-expression analyses revealed that canalized lncRNAs are functionally linked to canalized protein-coding genes, which are crucial regulators for caste differentiation. These canalized lncRNAs show striking tissue-specific enrichment consistent with canalized protein-coding genes. Functional validation through RNA interference revealed that canalized lncRNAs directly regulate caste-specific traits. Furthermore, juvenile hormone treatment capable of redirecting worker lncRNA expression profiles toward gyne-like patterns, with similar expression changes and tissue specificity to JH-responsive protein-coding genes, linking lncRNA regulation to established hormonal pathways controlling caste fate. Our findings establish lncRNAs as active architects of caste differentiation in social insects, demonstrating that these rapidly-evolving regulatory molecules contribute to the evolution and maintenance of social phenotypes through tissue-specific regulation of caste-associated developmental programs.

## Introduction

The evolution of social insect societies represents one of the major transitions in the history of life on Earth, fundamentally reshaping biological organization from individuals to superorganism (Szathmáry and Smith 1995). Ants provide an exceptional system for dissecting the general developmental and evolutionary mechanisms underlying this transition. Within ant colonies, genetically identical individuals follow divergent developmental trajectories that lead to the formation of morphologically and functionally distinct castes. Queens typically develop wings, enlarged ovaries, and specialized reproductive physiology, while workers generally possess reduced reproductive organs, enhanced sensory systems, and behavioral repertoires optimized for colony maintenance (Hölldobler and Wilson 1990; Rajakumar et al. 2024). This remarkable developmental plasticity, where a single genome generates multiple discrete phenotypes with division of reproductive labor, parallels cell differentiation in multicellular organisms where somatic cells sacrifice their reproductive potential to support germline cells (Wheeler 1910; Boomsma and Gawne 2018). Both represent major evolutionary transitions characterized by reproductive division of labor and offer unique insights into how complex biological systems evolve and are maintained through epigenetically regulated developmental programs (Bonasio 2012; Sieber et al. 2021; Okwaro and Korb 2023).

Recent advances have begun to illuminate the molecular architecture of caste differentiation, revealing that juvenile hormone (JH) signaling as a central regulatory pathway capable of redirecting caste fate and inducing transcriptomic changes that parallel natural caste differentiation (Penick et al. 2012; Gospocic et al. 2021; Li et al. 2024). And MAPK and insulin pathway are another two crucial signaling pathways to mediate caste differentiation in ants (Vizueta et al. 2025). Comparative transcriptomic analyses have identified key protein-coding genes with canalized expression pattern, which was characterized by progressively increasing between caste differences and decreasing within caste variation during development, that directly mediate caste-specific trait formation (Qiu et al. 2022). This concept of expression canalization extends Waddington’s phenotypic canalization framework to the molecular level (Waddington 1942; Waddington 1957), providing a powerful approach for identifying functionally relevant developmental regulators. However, protein-coding genes represent only one component of the regulatory landscape. Long non-coding RNAs (lncRNAs), transcripts longer than 200 nucleotides that lack protein-coding potential yet profoundly influence gene expression through diverse mechanisms including chromatin modification, transcriptional interference, and post-transcriptional regulation (Kapusta et al. 2013; Quinn and Chang 2016; Fernandes et al. 2019; Statello et al. 2021; Mattick et al. 2023), add a critical but largely unexplored regulatory layer to developmental systems.

Through mechanisms such as *cis-* or *trans-*interactions, modulation of chromatin status, and interactions with miRNAs, lncRNAs can profoundly influence gene expression and contribute to phenotypic outcomes (Ponting et al. 2009; Kopp and Mendell 2018; Statello et al. 2021). For instance, the lncRNA *PAHAL*, transcribed from the antisense strand of the *PAH* gene, modulates behavioral plasticity in locust by interacting with *PAH* promoter in *cis*, thereby modulating dopamine biosynthesis (Zhang et al. 2020). In butterfly, the lncRNA *ivory* controls the wing coloration patterning (Livraghi et al. 2024). *ivory* also modulates the intraspecific melanic wing color polymorphisms by *cis*-regulatory elements in buckeye butterfly (Fandino et al. 2024).

Among social insects, lncRNAs exhibit caste- or task-specific expression patterns in different lineages, including honeybees (Chen and Shi 2020), ants (Shields et al. 2018; Gao et al. 2020), wasps (Ferreira et al. 2013), and bumblebees (Collins et al. 2021). Despite accumulating evidence supporting the regulatory roles of lncRNAs, their functional contributions to caste differentiation remain enigmatic. Several fundamental questions persist: Do lncRNAs merely reflect developmental states passively, or do they actively regulate caste-specific developmental programs? If regulatory, do they exhibit canalized expression patterns similar to functionally important protein-coding genes as they share common transcriptional machinery? How are lncRNA expression profiles integrated with established hormonal pathways controlling caste determination? Are lncRNAs preferentially expressed in tissues critical for caste-associated functions? The arbitrary definition of lncRNAs based solely on length and absence of coding potential, combined with their rapid sequence evolution and diverse regulatory mechanisms, has impeded systematic functional investigation (Statello et al. 2021; Mattick et al. 2023). Consequently, whether lncRNAs contribute meaningfully to the evolution and maintenance of superorganism, or represent transcriptional noise, remains unresolved, leaving a significant gap in our understanding of the complete molecular architecture underlying one of biology’s major evolutionary transitions.

In this study, we address these questions through comprehensive transcriptomic analyses across complete developmental trajectories in two evolutionarily and ecologically distinct ant species, *Monomorium pharaonis* and *Acromyrmex echinatior* . By applying the canalization framework previously established for protein-coding genes, we systematically characterized lncRNA expression dynamics across development and test whether canalized lncRNAs are functionally involved in producing caste-specific phenotypes. Through functional experiments targeting candidate lncRNAs, we provide direct evidence for their regulatory roles in caste development. We further investigate how lncRNA regulation interfaces with juvenile hormone signaling. Our results establish lncRNAs as active regulators of caste development in social insects and provide mechanistic insights into how epigenetic factors contribute to the evolution and maintenance of complex social phenotypes.

## Results

### Characterization of lncRNAs in ants

To investigate the biological function of lncRNAs in ant caste development, we conducted comprehensive transcriptomic analysis across complete developmental trajectories in two ecologically and evolutionary distinct ant species. Our dataset encompassed 16 developmental stages spanning from embryo to adult in *M. pharaonis,* and 8 stages from larva to adult in *A. echinatior* (Qiu et al. 2022). These two species probably represent typical modes of colony and caste differentiation across most extant ants: *M. pharaonis* is a polygynous (multiqueen) species with monomorphic worker caste, and their caste fate is specified blastogenically (Khila and Abouheif 2010; Pontieri and Linksvayer 2021), while in *A. echinatior*, its colony always only has a single queen and polymorphic workers, and their caste determination occurs during early larval development (Bekkevold et al. 1999; Adams et al. 2021). By leveraging high-quality genome assemblies generated by Global Ant Genomics Alliance project (Boomsma et al. 2017) and our comprehensive RNA-seq dataset, we reconstructed the transcript models for both species and identified 10400 and 10772 lncRNAs in *M. pharaonis* and *A. echinatior*, respectively, after stringent filtering to remove transcripts shorter than 200 nt or with protein-coding potential (Materials and Methods; *SI Appendix*, Fig. S1).

Our comparative analysis revealed that ant lncRNAs exhibit distinctive sequence characteristics that distinguish them from protein-coding genes. Most notably, lncRNAs demonstrated remarkably rapid sequence evolution, as evidenced by substantially fewer orthologous lncRNAs between the two species compared to protein-coding genes (*SI Appendix*, Fig. S2*A*). This evolutionary lability appears to be driven, at least in part, by the significantly higher proportion of transposable element (TE) insertions within lncRNA sequences compared to protein-coding genes in *M. pharaonis* (*SI Appendix*, Fig. S2*B*). This finding highlights the important role of TEs in both the origin and ongoing diversification of lncRNA repertoires. Additionally, lncRNAs exhibited simpler structural organization, containing fewer exons than protein-coding genes in both species (*SI Appendix*, Fig. S2*C*), consistent with patterns observed across diverse taxa and suggesting less conservation and functional constraint in architecture of lncRNAs.

The developmental expression dynamics of lncRNAs in *M. pharaonis* revealed intriguing temporal patterns that align with key developmental transitions. Following the zygotic genome activation at 24 hours post-fertilization (Rajakumar et al. 2024), stable lncRNA numbers were detected (TPM>1) within the same developmental stages (embryos, larvae, pupae and imago), while the developmental transitions are often accompanied by dramatic changes in the number of expressed lncRNAs (Fig. 1*A*). These temporal dynamics coincide with the expression dynamics of protein-coding genes (*SI Appendix*, Fig. S3*A*), implying rapid morphological remodeling during these transition stages (Du et al. 2023; Wilkens et al. 2025).

**Fig. 1.**
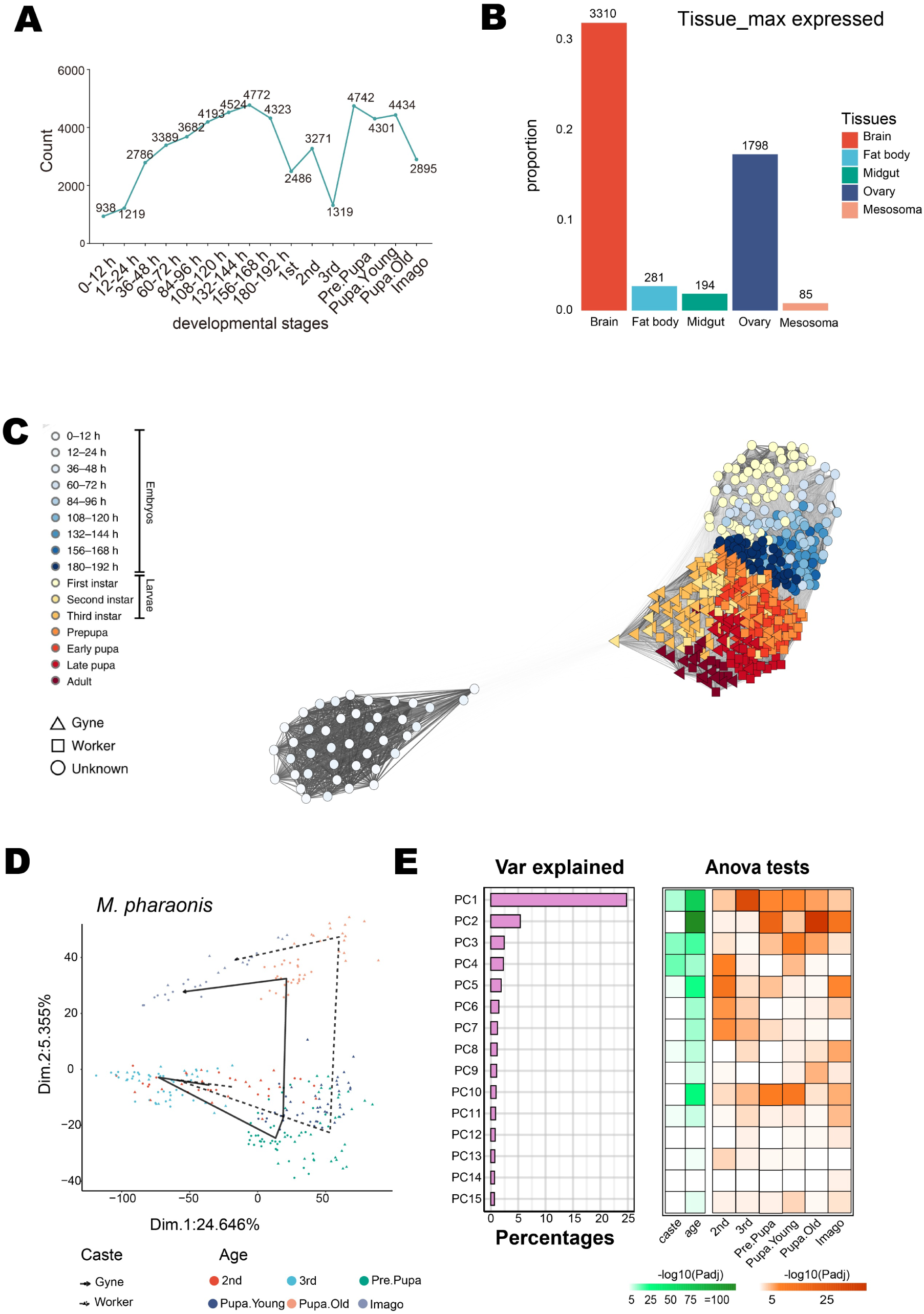
Developmental lncRNA atlases. (*A*) Number of expressed lncRNAs (the average TPM > 1 in all samples) in each developmental stage in *M. pharaonis*. Notably, castes can be distinguished since the 2^nd^ instar larval stage, the count includes lncRNAs from both castes (worker and gyne). (*B*) Number of lncRNAs with maximum expression in each tissue type. RNA-seq data from 5 tissues of imago gynes (within 3 days) were used to calculate expression levels. Maximum expression refers to having the highest expression level in the samples of this tissue compared to other tissues. (*C*) *M. pharaonis* developmental trajectories reconstructed from lncRNA profiles of 502 individuals covering all developmental stages. Individuals with similar lncRNA profiles cluster together based on Spearman’s correlation coefficient. Shading of connecting lines reflects the correlation of mutual lncRNA profiles. Unknown, including embryonic and 1^st^ instar larval stages, means the caste of individual cannot be distinguished morphologically. (*D*) The developmentally dynamic changes of lncRNA in *M. pharaonis* are illustrated by Principal component analysis, from the morphologically distinct larval stage (2^nd^ instar) to adults. The lines with arrows show the direction of development. The colors show the developmental stages. The lines of solid and dotted indicate gyne and worker, respectively. (*E*) For the left panel, percentage of variation explained by the first 15 principal components from the PCA described in (*D*). For the middle panel with green color, we tested if there was a significant difference between castes or developmental stages for each principal component. For the right panel with orange color, we tested if there was a significant difference between any query group (developmental stage) versus all other collapsed groups. All tests were performed with the aov function in R.

To address the potential functionality of lncRNAs, we examined their tissue-specific expression across five adult gyne organs (brain, mesosoma, fat body, midgut and ovary from newly eclosed gynes within 3 days) and four worker organs (brain, mesosoma, fat body and midgut from young workers) in *M. pharaonis*. Both gyne and worker exhibit a striking expression bias toward neural tissues of lncRNAs. More than half of all detected lncRNAs exhibited highest expression in the brain (Fig. 1*B* and *SI Appendix*, Fig. S3*B*), a pattern consistent with observations in *Harpegnathos saltator*, another ant species (Shields et al. 2018). This pronounced neural enrichment suggests that lncRNAs may play particularly important regulatory roles in neural development and potentially in the evolution of complex behaviors that characterize social insects. Compared with protein-coding genes, lncRNAs exhibit higher tissue specificity and lower expression level across developmental stages (*SI Appendix*, Fig. S3*C* and *D*), suggesting potential tissue-specific functions.

To assess the relationship between lncRNA and caste development, we reconstructed the developmental trajectories using lncRNA expression profiles. In *M. pharaonis*, maternal lncRNAs expressed during the earliest stages (0-24 h) clustered distinctly from those expressed after zygotic genome activation, indicating functional distinct roles for maternally inherited versus zygotically expressed lncRNAs. From 24 hours onward, lncRNA profiles showed continuous developmental progression, with adjacent stages clustering together while maintaining stage-specific signature (Fig. 1*C*). Principal component analysis (PCA) further revealed that the first two principal components captured variation driven by both developmental stage and caste identity (Fig. 1*D* and *E*), strongly suggesting that lncRNAs contribute to both general developmental process and caste-specific differentiation programs. Similarly, in *A. echinatior*, expression profiles clustered by both developmental stage and caste identity (*SI Appendix*, Fig. S4*A*), similarity matrices confirming discrete profiles for each developmental stage and caste combination (*SI Appendix*, Fig. S4*B*). This dual role was reinforced by our identification of differentially expressed lncRNAs (DElncs) between castes in both species (*SI Appendix*, Fig. S4*C*), which increased substantially during metamorphosis, when caste-specific phenotypes become morphologically apparent. These results collectively demonstrate that the expression of lncRNAs likely contribute to caste differentiation during developmental process, positioning them as potential molecular players in the evolution of superorganisms.

### Canalized lncRNAs enriched in caste-associated organs

To comprehensively understand how lncRNAs contribute to the caste-specific developmental programs, we focused our analysis on the phenomenon of expression canalization—characterized by continuously increasing transcriptomic difference between castes coupled with decreasing transcriptomic variation within each caste as development proceeds. Our previous work demonstrated that protein-coding genes exhibiting canalized expression patterns are intimately associated with caste differentiation (Qiu et al. 2022), and we hypothesized that lncRNAs might follow similar regulatory logic as they share the similar biogenesis process.

To quantify expression tendency of lncRNA profiles, we calculated the developmental potential of each individual based on their lncRNA expression profiles, measuring this as the deviation from the average target caste profiles (gyne/worker) in the subsequent development stages (Materials and Methods). Individuals with a closer distance to the gyne profiles have positive values, reflecting gyne-biased developmental potential, whereas negative values indicate worker-biased trajectories. Remarkably, we found the overall lncRNA expression exhibited pronounced canalization patterns mirroring those previously documented for protein-coding genes (Qiu et al. 2022). In both *M. pharaonis* and *A. echinatior,* individuals from different castes showed overlapping developmental potential during early development, but gradually diverged into distinct, caste-specific clusters as development progressed (Fig. 2*A* and *SI Appendix*, Fig. S5*A*). This divergence was accompanied by progressively decreasing within-caste variation in lncRNA profiles, confirming transcriptomic canalization at the lncRNA level (*SI Appendix*, Fig. S5*B*).

**Fig. 2.**
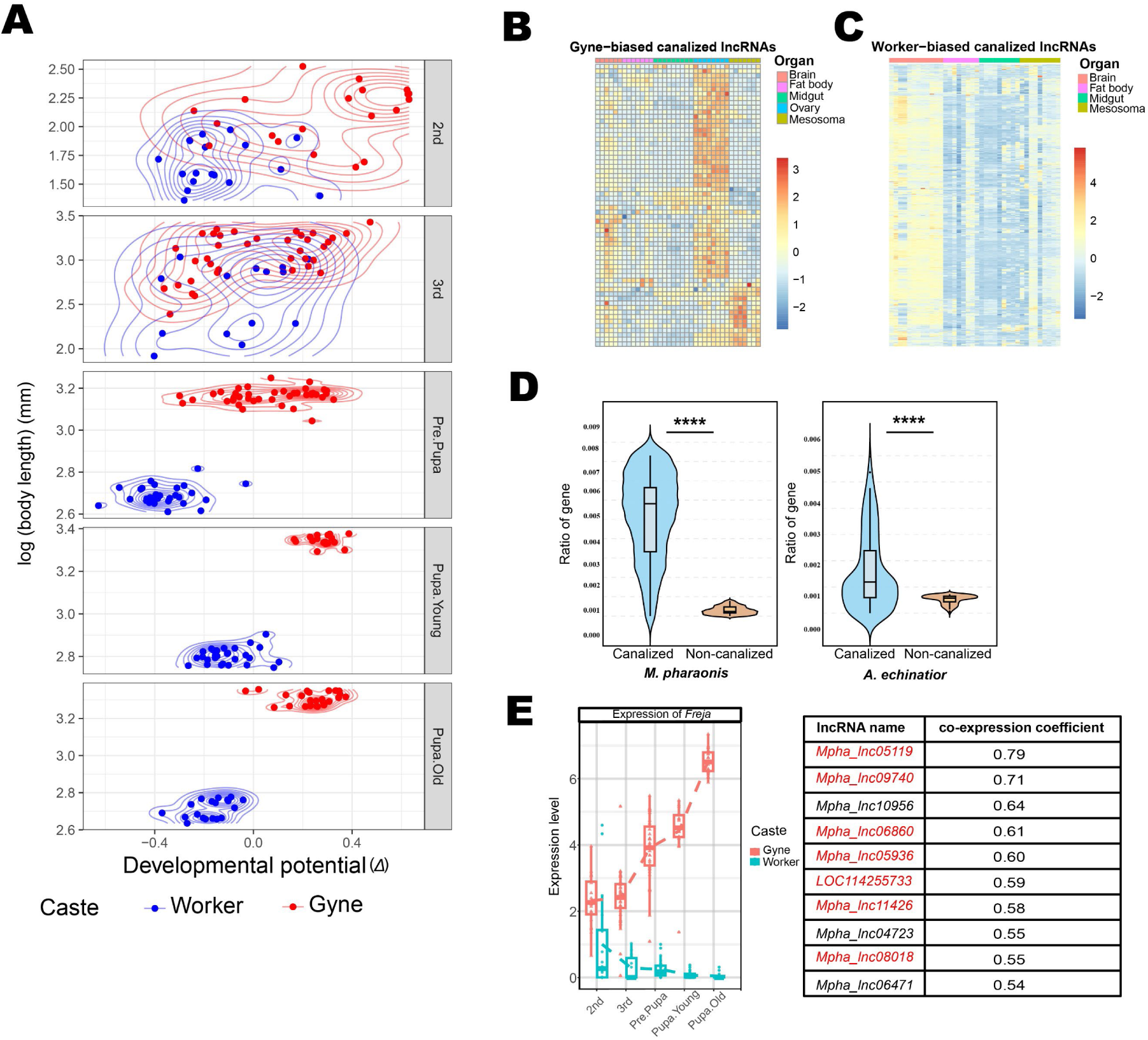
Individual lncRNA profiles indicate their caste canalization and their co-expression with canalized protein-coding genes. (*A*) Developmental potential (*Δ*) scores for individual gynes and workers in *M. pharaonis*. The *Δ* score reflects the difference in lncRNA profile distances between target individuals and an average gyne and worker lncRNA profiles in the next developmental stage, reflecting the developmental tendency towards corresponding caste phenotype. A positive value means the lncRNA profile has a gyne-biased developmental potential. Conversely, a negative one means a worker-biased developmental potential. (*B*) Tissue-specific relative expression of gyne-biased canalized lncRNAs in *M. pharaonis*. Heatmap shows expression value (TPM) for each lncRNA by row, with columns clustered by tissue and rows clustered by expression similarity. The expression values are normalized and shown as Z-scores. 5 gyne tissues, brain, fat body, midgut, ovary and mesosoma, were used for gyne-biased canalized lncRNAs. The lncRNA list was formed according to canalization score (n = 62, C.score > 1). (*C*) Tissue-specific relative expression of worker-biased canalized lncRNAs in *M. pharaonis*. Heatmap shows expression value (TPM) for each lncRNA by row, with columns clustered by tissue and rows clustered by expression similarity. The expression values are normalized and shown as Z-scores. 4 worker tissues, brain, fat body, midgut and mesosoma, were used for worker-biased canalized lncRNAs. The lncRNA list was formed according to canalization score (n = 761, C.score < -1). (*D*) Co-expression between canalized lncRNAs and protein-coding genes in both species. For each canalized lncRNA, the number of canalized protein-coding genes among its top 10 co-expression genes was counted and normalized by the total gene number. The number of canalized lncRNAs was 823 in *M. pharaonis* and 185 in *A. echinatior*. Comparisons were made using Wilcoxon test (*M. pharaonis*, pvalue < 2.2e-16; *A. echinatior*, pvalue = 6.375e-15; two-tailed t-test; **** pvalue < 0.0001). (E) The protein-coding gene with most significant canalized expression pattern, *Freja* (Qiu et al. 2022), and its top 10 co-expressed lncRNAs. The left panel shows the expression pattern of *Freja*; the right table represents the co-expression coefficient between these lncRNAs and *Freja*. Canalized lncRNAs are shown in red, and non-canalized lncRNAs are shown in black.

Then, we identified lncRNAs with canalized pattern using the same method as our previous work via the ratio of between-caste lncRNA expression difference and stage-specific expression variance within castes (Qiu et al. 2022). Considering the low expression level and the high tissue specificity of lncRNAs, the stringent canalization scoring criteria for protein-coding genes will filter out some lncRNAs. Applying a criteria of |Canalization score|(C.score) ≥ 1, we identified 832 canalized lncRNAs in *M. pharaonis* (gynes versus workers), of which 62 are gyne-biased and 761 are worker-biased (Dataset S1). To assess the functional relevance of these canalized lncRNAs, we examined their tissue-specific expression patterns using adult tissue RNA-seq in *M. pharaonis*. The results revealed the lncRNAs with canalization patterns exhibit tissue-specific enrichment in organs critical for caste-associated functions. Gyne-biased canalized lncRNAs exhibited remarkable tissue specificity, with more than half showing significantly elevated expression in the ovary (Hypergeometic test, pvalue = 7.76e-12)(Fig. 2*B* and *SI Appendix*, Fig. S6*A*), the primary reproductive organ that distinguishes queens from sterile workers. Additionally, several gyne-biased lncRNAs were enriched in the mesosoma, the body segment housing flight muscles and other structures essential for the reproductive dispersal flights that characterize queen behavior (Hypergeometic test, pvalue = 1.38e-16)(Fig. 2*B* and *SI Appendix*, Fig. S6*A*). Although gynes in *M. pharaonis* are flightless, our previous work revealed that they possess more developed flight muscle than workers (Li et al. 2024). In contrast, very few gyne-biased lncRNAs showed enrichment in the midgut, brain, or fat body, suggesting that the canalized lncRNAs in gynes are selectively associated with reproductively relevant rather than housekeeping functions. While the worker-biased canalized lncRNAs displayed an even more pronounced tissue bias, with nearly all showing enrichment in the brain (Hypergeometic test, pvalue = 1.7e-233)(Fig. 2*C* and *SI Appendix*, Fig. S6*A*). This neural enrichment is consistent with the characteristic of worker caste, such as cognitive and behavioral specializations that support complex foraging behaviors, nest maintenance, and brood care. This tissue distribution pattern of canalized lncRNAs exhibits strong concordance with canalized protein-coding genes, which are known to regulate caste differentiation, suggesting comparable functional significance of canalized lncRNAs (*SI Appendix*, Fig. S6*B* and *C*).

To assess the potential functions of canalized lncRNAs in development of caste-associated traits, we analyzed their co-expression patterns with protein-coding genes—a widely used strategy for inferring lncRNA function (Liao et al. 2011; Sarropoulos et al. 2019). Using Spearman correlation analysis, we discovered that lncRNAs with canalized expression patterns were significantly more likely to co-express with protein-coding genes that also exhibited canalized patterns in both *M. pharaonis* and *A. echinatior* (Fig. 2*D*). This co-expression network analysis revealed functional clustering around key developmental regulators. Most notably, *Freja*, a key regulator of queen phenotypes that we previously identified (Qiu et al. 2022), showed strong co-expression with multiple canalized lncRNAs (Fig. 2*E*). Of the 10 lncRNAs most strongly co-expressed with *Freja*, seven showed canalized expression, all of which were enriched in ovarian tissue, mirroring *Freja*’s own pattern (*SI Appendix*, Fig. S7). This convergent expression pattern suggests that canalized lncRNAs may function as components of larger regulatory modules centered around master developmental regulators.

### Canalized lncRNA regulates caste-related phenotypes

The identification of canalized lncRNAs with tissue-specific expression patterns and co-expression with master developmental regulators provided strong circumstantial evidence for their functional roles in caste differentiation. To demonstrate the function of canalized lncRNAs in caste-associated trait development, we functionally validated representative lncRNAs through RNA interference (RNAi) experiment. Our selection criteria prioritized lncRNAs that exhibited strong canalization patterns, tissue-specific enrichment in caste-relevant organs, and potential mechanistic links to known developmental pathways.

Our candidate, designated *ASfln* (C.score = 2.57, pvalue = 0.021), transcribes from the antisense strand of *flightin* (C.score = 9.82, pvalue = 0.009)(Fig. 3*A*), a gene encoding a crucial structural protein required for proper flight muscle development and function (Vigoreaux et al. 1993). RNA-seq and RT-qPCR data showed that both *ASfln* and *flightin* are expressed in flight muscle and exhibit similar temporal expression dynamics (Fig. 3*B*, *SI Appendix*, Fig. S8*A* and *B*). The genomic organization of *ASfln* and *flightin* suggests potential for direct regulatory interactions, as antisense lncRNAs frequently modulate the expression of their sense-strand partners through various mechanisms including chromatin modification, transcriptional interference, or post-transcriptional regulation (Ponting et al. 2009; Kopp and Mendell 2018; Statello et al. 2021). Flight capability represents one fundamental caste distinction in ants, as reproductive individuals typically possess well-developed flight muscles, while workers generally lack functional flight apparatus (Hölldobler and Wilson 1990; Abouheif and Wray 2002). The coordinated expression of *ASfln* and *flightin* throughout development, coupled with their shared enrichment in flight muscle tissue, suggested that *ASfln* might function as a *cis*-regulatory element controlling *flightin* expression and, consequently, flight muscle development.

**Fig. 3.**
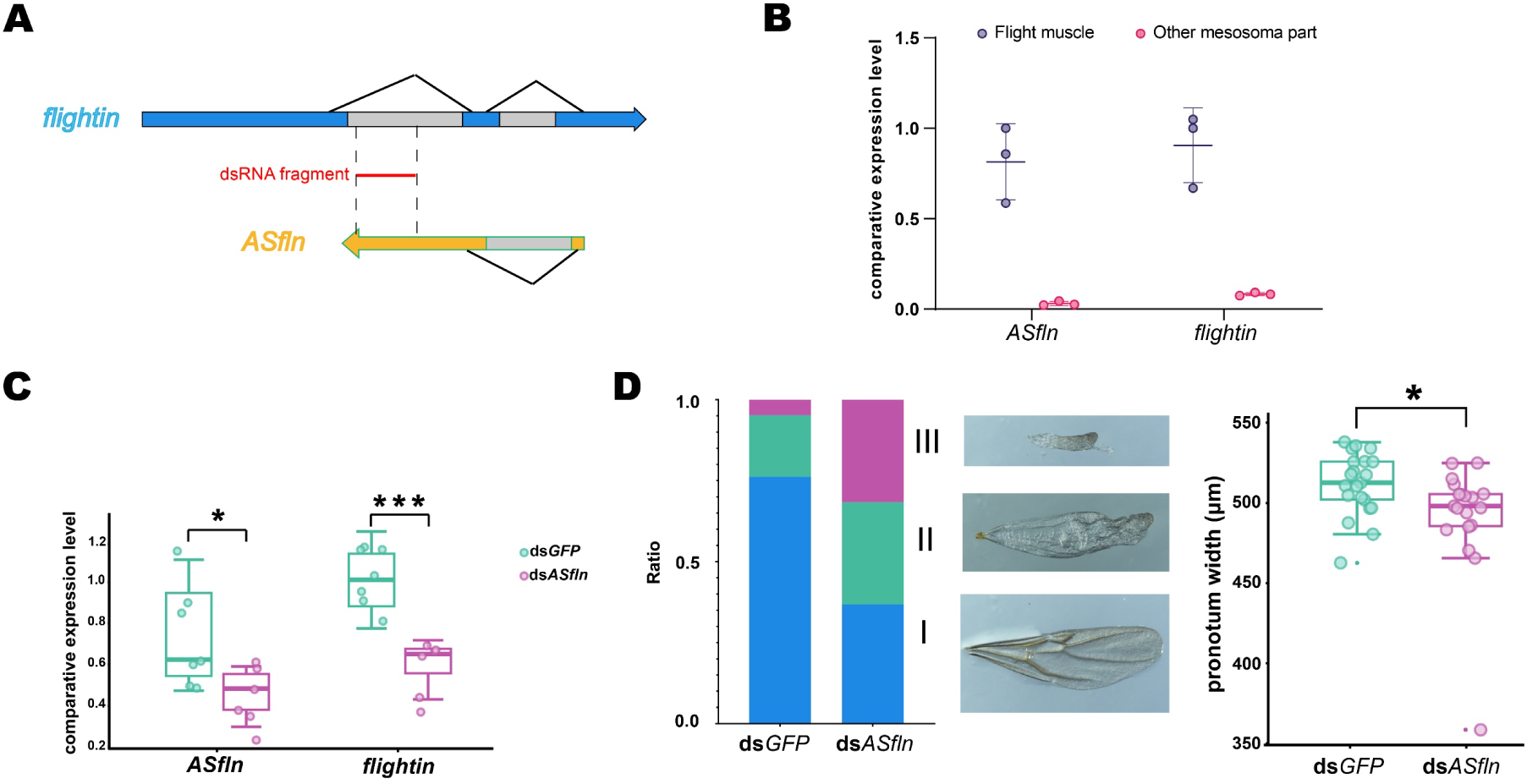
Expression and functional validation of lncRNA with canalization pattern. (*A*) The genomic locations of *flightin* and *ASfln*. Arrows indicate their transcription directions. The grey boxes represent introns; blue and yellow boxes represent exons of *flightin* and *ASfln*, respectively. Red solid line between *flightin* and *ASfln* indicates dsRNA fragment used for specific repressing the expression level of *ASfln*. The dotted lines indicate the position of the dsRNA in the gene structures of *flightin* and *ASfln*. (*B*) The expression localization of *ASfln* and *flightin* in mesosoma. The mesosoma was dissected into flight muscle and other parts. Then the expression levels were detected by RT-qPCR. Three biological replicates were used (mean ± SD). (*C*) The comparative expression level of *ASfln* and *flightin* when *ASfln* is knocked down with dsRNA. After 48h of dsRNA injection, the expression levels of *ASfln* and *flightin* were investigated by RT-qPCR. pvalue = 0.0324 for *ASfln* (two-tailed t-test; n = 7; * pvalue < 0.05); pvalue = 0.0005 for *flightin* (two-tailed t-test; n = 6; ***pvalue < 0.001). (*D*) RNAi phenotype. The wing development and pronotum width were recorded and measured on the 1^st^ day of adult gyne. The left histogram categorizes wing type: blue (type I) for normal wings, green (type II) for wings with folded distal area, and purple (type III) for wholly defective wings. The ds*ASfln* group exhibited significantly more defects (Chi-square test, pvalue = 0.0246). The right boxplot shows the comparison of pronotum width between two groups. pvalue = 0.0374 (two-tailed t-test, n = 21 for ds*GFP* and n = 19 for ds*ASfln*; * pvalue < 0.05).

RNA interference targeting *ASfln* during the critical larval and pupal stages, when wing and flight muscle development occurs, resulted in increased incidence of wing developmental defects in emerging gynes. These malformations ranged from subtle asymmetries to more severe structural abnormalities that would likely compromise flight performance (Chi-square test, pvalue = 0.0246)(Fig. 3*D*). This phenotype is similar to the wing defect caused by *flightin* knockdown in other species (Chang et al. 2022). Additionally, morphometric analysis revealed significant reduction in pronotum width of *ASfln* RNAi group, a measurement that reflects the overall development of the thoracic region housing flight muscle (pvalue = 0.0374)(Fig. 3*D* and *SI Appendix*, Fig. S8*C*). Of note, we observed a significant decreased expression of *flightin* following *ASfln* knockdown (pvalue = 0.0005)(Fig. 3*C*). This regulatory relationship appears to operate through *cis*-acting mechanisms, as the dsRNA fragment used for *ASfln* targeting was specifically designed to avoid direct interference with *flightin* transcripts (Fig. 3*A*). The coordinate reduction in both *ASfln* and *flightin* expression following antisense lncRNA suppression suggests that *ASfln* functions as a positive regulator of *flightin* transcription, possibly through chromatin modifications or recruitment of transcriptional machinery to the *flightin* locus.

Reproductive division of labor is a hallmark of eusociality in ants (Khila and Abouheif 2008; Khila and Abouheif 2010). However, the underlying molecular mechanisms of reproductive regulation are yet to be comprehensive resolved. To figure out whether lncRNAs participate in this process, an ovary-enriched lncRNA (*lncov*)(C.score = 1.86, pvalue = 0.136) was selected for functional validation. This lncRNA displays a pronounced canalized expression pattern with progressive gyne-specific upregulation throughout development and demonstrates striking ovary-specific enrichment in adults (*SI Appendix*, Fig. S9*A*). Remarkably, the spatial expression pattern of *lncov* suggests a potential role in oogenesis, a process that represents one of the most fundamental distinctions between reproductive and sterile castes in ant societies (*SI Appendix*, Fig. S9*B*) (Wang et al. 1994; Khila and Abouheif 2008; Khila and Abouheif 2010), Given *lncov*’s expression pattern, we hypothesized that this lncRNA might function as a key regulator of gyne-specific reproductive development. To test this hypothesis, we performed targeted knockdown experiments using RNAi in adult gynes. Following *lncov* supression, treated gynes exhibited significant reductions in both oocyte number and size compared to control individuals (*SI Appendix*, Fig. S9*C* and *D*). The specificity of this phenotype, combined with *lncov*’s restricted expression pattern, strongly supports a direct regulatory role of *lncov* in gyne reproductive development.

### Juvenile hormone signaling mediates the expression of lncRNAs mirroring that of protein-coding genes

Juvenile hormone (JH) is a key regulator for caste differentiation across diverse social insects. In *M. pharaonis*, JH treatment during larval stages can dramatically redirect worker development toward gyne-like phenotypes, inducing both morphological and transcriptomic changes that mirror natural gyne development (Li et al. 2024). However, the relationship between lncRNAs expression and JH signaling during caste development is still to be investigated.

In our previous work, we treated *M. pharaonis* worker with JH during the early 3^rd^ instar larval stage and collected samples for RNA-seq across five critical developmental time points spanning the 3^rd^ instar larval, prepupal, and young pupal stages (Li et al. 2024)—the developmental window during which caste fate becomes irreversibly fixed. Here, we compared the lncRNA profiles of gynes, workers, and JH-treated workers (hereafter “JH workers”) using these whole-body RNA-seq data. We calculated caste tendency scores based on spearman correlation coefficients to quantify the similarity between JH-treated workers and natural caste profiles. The results revealed a striking developmental reprogramming of lncRNA expression following JH treatment. As development progressed, the lncRNA expression profiles of JH workers shifted progressively towards that of natural gynes (Fig. 4*A*), indicating that JH can fundamentally reprogram lncRNA expression in a caste-specific manner. This temporal shift was not merely a static response but rather a dynamic developmental redirection that became increasingly pronounced with time, suggesting that JH initiates cascading regulatory changes that amplify throughout development.

**Fig. 4.**
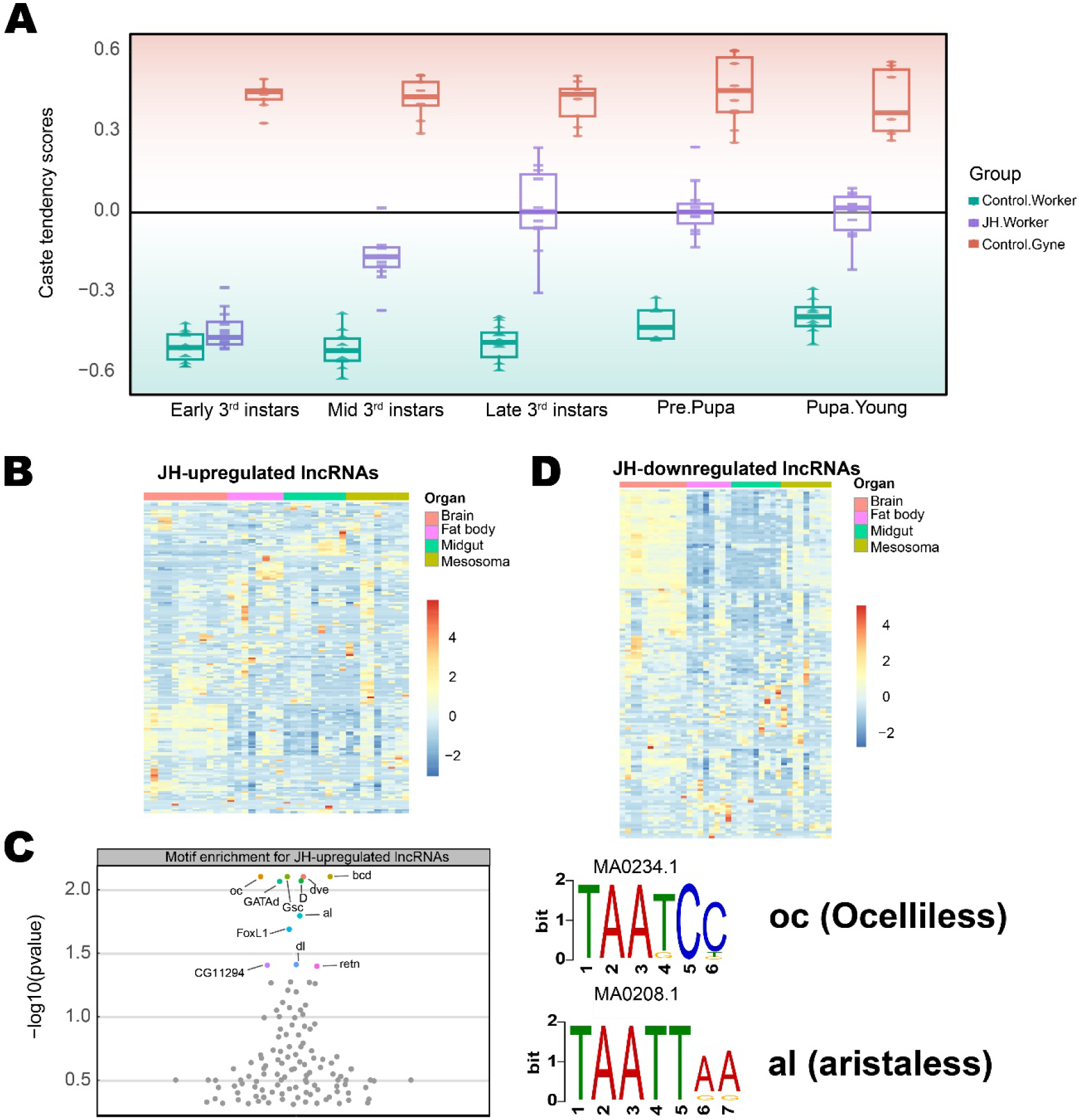
Juvenile hormone signaling mediates lncRNA expression. (*A*) Overall changes of lncRNA profiles in JH workers. Caste tendency scores (CTS) of individuals indicate the similarities between a JH worker and the average same-age gyne and worker, calculating according to Euclidean distance. See methods for the formula. (*B*) Tissue-specific relative expression of lncRNAs up-regulated in JH workers in *M. pharaonis*. Heatmap shows expression value (TPM) for each lncRNA by row, with columns clustered by tissue and rows clustered by expression similarity. The expression values are normalized and shown as Z-scores. 4 worker tissues, brain, fat body, midgut and mesosoma, were used. This lncRNA list was formed according to *cd* score (n = 168, *cd* > 1, pvalue ≤ 0.05). (*C*) Motif enrichment for the regulatory regions of lncRNAs up-regulated by JH. The regulatory region was defined as 2.5 kb within the upstream and downstream of TSS. The insect transcription factor binding site (TFBS) motifs from JASPAR database were used as query. All TFs with pvalue < 0.5 were presented. TFs with pvalue < 0.05 were presented with a colored dot, others were grey. The sequence logo of two enriched motifs is shown on the right. (*D*) Tissue-specific relative expression of lncRNAs down-regulated in JH workers in *M. pharaonis*. Heatmap shows expression value (TPM) for each lncRNA by row, with columns clustered by tissue and rows clustered by expression similarity. The expression values are normalized and shown as Z-scores. 4 worker tissues, brain, fat body, midgut and mesosoma, were used. This lncRNA list was formed according to *cd* score (n = 149, *cd* < -1, pvalue ≤ 0.05).

To better understand this dynamic developmental process, we first identified differentially expressed lncRNAs between control workers and gynes (caste-DElncs) and between control workers and JH workers (JH treatment DElncs) for each time point (Materials and Methods)(Dataset S2). The significant overlap between caste-DElncs and JH treatment DElncs confirms that the expression levels of lncRNAs in JH workers are shifted towards to that of gynes (*SI Appendix*, Fig. S10*A* and *C*). This result aligns with the changes of protein-coding genes (*SI Appendix*, Fig. S10*B*-*D*). We then focus on the tissue distribution of these overlapping DElncs (oDElncs) and DEGs (oDEGs). Both (upregulated/downregulated) oDElncs and oDEGs are likely enriched in the similar tissues in each time point (*SI Appendix*, Fig. S10*E*), hinting their functional relevance.

To further identify specific lncRNAs responsive to JH signaling (JH-responsive lncRNAs), we adapted our previously developed canalization-disrupted score (*cd* score) (Li et al. 2024), which quantifies the extent to which JH treatment perturbs canalized gene expression patterns (Materials and Methods). Applying a stringent threshold of |*cd*| > 1 and pvalue < 0.05, we identified 168 gyne-biased lncRNAs that were upregulated by JH treatment and 149 worker-biased lncRNAs that were downregulated (Dataset S3). Notably, among the top 10 lncRNAs, the directional changes were consistent: gyne-biased lncRNAs were invariably upregulated while worker-biased lncRNAs were consistently downregulated across most time points (*SI Appendix*, Fig. S11*A* and *B*).

The tissue-specific expression patterns of these JH-responsive lncRNAs provided crucial insights into their potential functional roles. We examined the expression level of these JH-responsive lncRNAs across the 4 worker tissues and compared with JH-responsive protein-coding genes. Both the lncRNAs and protein-coding genes that were downregulated by JH show higher expression in the brain than other tissues (Fig. 4*D* and *SI Appendix*, Fig. S11*C*), suggesting their potential contribution to the elaborate brain structures and complicated behaviors in the workers. For the JH-upregulated lncRNAs and protein-coding genes, they do not show obvious tissue specificity (Fig. 4B and *SI Appendix*, Fig. S11*D*). This result is consistent with the phenotypic changes that multiple tissues were remodeled in parallel in JH workers (Li et al. 2024). To elucidate the molecular mechanisms underlying JH-mediated lncRNA regulation, we conducted transcription factor binding site analysis in the putative promoter regions of JH-upregulated lncRNAs because promoter region is the most conserved region and regulates the expression of lncRNAs (Derrien et al. 2012; Mattick et al. 2023). Using insect transcription factor binding site (TFBS) motifs from the JASPAR database as query (Rauluseviciute et al. 2024), we identified significant enrichment of binding sites for 11 transcription factors in gyne-biased lncRNAs (pvalue ≤ 0.05)(Fig. 4*C*; Dataset S4). Eight of which are established regulators of developmental processes (Dataset S4), particularly in tissues relevant to caste differentiation. For instance, *ocelliless* (*oc*) is an essential TF for ocelli development. Its loss eliminates ocelli in *Drosophila* (Jean-Guillaume and Kumar 2022). Another highly enriched TF, *aristaless* (*al*), is a conserved regulator of imaginal disc patterning in invertebrates that has been implicated in wing development across diverse insect species including *Drosophila* (Campbell and Tomlinson 1998), *Junonia* (Martin and Reed 2010) and *Heliconius* (Bayala et al. 2023).

Take together, our finding reveals that JH-responsive lncRNAs show a similar expression dynamics and tissue-specificity with the JH-responsive protein-coding genes. This result suggests that JH-mediated caste differentiation operates through multiple molecular layers, with lncRNAs representing a previously unrecognized but crucial component of this regulatory architecture. The identification of specific transcription factors enriched in JH-upregulated lncRNA promoters provides a mechanistic framework for understanding how hormonal signals are translated into caste-specific lncRNA expression programs, revealing lncRNAs as both targets and potential mediators of caste development in social insects.

## Discussion

### lncRNAs as active regulators of caste development in social systems

Caste differentiation in ants is a fascinating topic for both developmental biology and evolutionary biology (Wheeler 1910; Waddington 1957; Boomsma and Gawne 2018). As queens and workers in a colony share the identical genome, their divergent development are likely to be associated with epigenetic factors (Bonasio 2012; Bonasio 2014; Oldroyd and Yagound 2021; Orr and Goodisman 2023; Lebedev et al. 2024). Comparative studies have implicated several such mechanisms, including DNA methylation (Bonasio et al. 2012; Oldroyd and Yagound 2021), histone modification (Bonasio et al. 2010; Simola et al. 2013; Wojciechowski et al. 2018), RNA editing (Li et al. 2014), and non-coding RNAs (ncRNAs) (Shi et al. 2015; Shields et al. 2018; Gao et al. 2020) in caste differentiation. Our study provides the first demonstration that long non-coding RNAs function as active regulators of caste development in social insects. The identification of over 832 canalized lncRNAs in *M. pharaonis*, characterized by progressively increasing caste bias and decreasing within-caste variation throughout development, reveals a previously unrecognized layer of epigenetic regulation underlying caste differentiation. Critically, our functional validation experiments with *lncov* and *ASfln* demonstrate that these canalized expression patterns translate into direct regulatory control over caste-specific traits, rather than merely reflecting passive consequences of developmental commitment. The tissue-specific enrichment patterns we observed, with gyne-biased canalized lncRNAs concentrated in reproductive and flight muscle tissues, while worker-biased canalized lncRNAs predominate in neural tissues, suggest that lncRNA-mediated canalization operates through spatially restricted regulatory modules that fine-tune the development of caste-associated organs. This spatial organization of canalized lncRNA expression provides a mechanistic explanation for how developmental systems can simultaneously maintain phenotypic robustness within castes while generating dramatic morphological differences between castes, a fundamental requirement for the evolution and maintenance of eusociality. As lncRNAs also play crucial roles in cell differentiation and development in multicullar organisms (Hu et al. 2012; Fatica and Bozzoni 2014; Luo et al. 2016), which inspire us to believe that the different major evolutionary transition events may share the similar regulatory mechanism.

### Integration of lncRNA regulation with established caste determination pathways

A particularly significant finding of our study is the demonstration that lncRNA expression profiles are directly responsive to juvenile hormone signaling, the master regulatory pathway controlling caste fate in social insects. In *M. pharaonis*, JH treatment during larval stages can redirect development toward a gyne-like phenotype at both morphological and transcriptomic levels (Li et al. 2024). In *H. saltator*, the JH-induced transcription factor Kr-h1 modulates caste identity by regulating distinct gene sets in different castes in response to hormone signaling (Gospocic et al. 2021). Our observation extended the repertoire of JH-responsive molecules to include lncRNAs. We show that JH treatment can redirect worker lncRNA expression profiles toward gyne-like patterns. Moreover, JH-responsive lncRNAs exhibit similar expression dynamics (the ratio of JH-responsive lncRNAs among caste-biased lncRNAs) and tissue specificity as JH-responsive protein-coding genes, thereby establishing lncRNAs as downstream effectors of hormonal caste determination mechanisms, rather than independent regulatory systems. The enrichment of developmental transcription factor binding sites in JH-responsive lncRNA promoters—including key regulators such as ocelliless and aristaless, which control eye and wing development, respectively—reveals how hormonal signals are translated into tissue-specific lncRNA expression changes that ultimately drive morphological differentiation. Furthermore, our co-expression analyses linking canalized lncRNAs to established caste regulators such as *Freja* demonstrate that lncRNAs are integrated into conserved gene regulatory networks rather than operating as isolated regulatory modules. This integration suggests that lncRNAs may serve as evolutionary tuning mechanisms that allow fine-scale modification of caste phenotypes without disrupting core developmental programs, potentially facilitating the rapid evolution of species-specific caste morphologies observed across ant lineages.

### Evolutionary implications of epigenetic regulation in phenotypic development

Both the canalized lncRNAs and JH-responsive lncRNAs parallel the corresponding protein-coding genes in their expression pattern, tissue-specificity, and co-expression, suggesting their potential involvement in common regulatory networks. With the discovery of additional regulatory elements such as pseudogenes (Qian et al. 2022), microRNAs (Bartel 2009), and highly conserved elements (HCE) (Seki et al. 2017), our understanding of the genotype-phenotype map has expanded beyond the framework of the central dogma. The evolution of protein-coding genes is much slow (Huang and Zhang 2014); thus, the great phenotypic diversity of life should be created by various regulatory elements to assist organisms to adapt to changing environments. The rapid sequence evolution of lncRNAs (Ponting and Haerty 2022; Mattick et al. 2023), evidenced by low orthology rates between our two investigated species and numerous studies, suggests that the specific molecular sequences of lncRNAs may be evolutionarily labile. However, the conservation of canalized expression patterns and functional roles indicate the regulatory logic and developmental functions can be maintained through selection on expression patterns rather than sequence conservation. This situation occurs in various species. For instance, defects in brain development caused by linc-*birc6* in zebrafish can be rescued by injecting its orthologs from humans or mice despite limited sequence similarity (Ulitsky et al. 2011). In ants, the ancient yet rapidly evolving lncRNA *ANTSR* has assumed a role in sex determination and this function may be evolved early in ant evolution (Pan et al. 2024). Overall, our findings provide a new molecule as a regulatory layer for understanding how complex phenotypes evolve, thereby advancing insights into the genotype-phenotype map and even the developmental mechanisms of major evolutionary transitions.

### From developmental canalization to molecular robustness

Waddington’s concept of canalization describes the stabilization of developmental trajectories against genetic variation or environmental perturbations, ensuring that organisms generate reliable, stereotyped phenotypes (Waddington 1942). This idea of developmental canalization has long attracted the attention of both developmental and evolutionary biologists, as it appears conceptually opposed to phenotypic plasticity. These two concepts deal with one of the most fundamental questions in biology: how the living organisms are shaped and diversified through development and evolution (Debat and Le Rouzic 2019). Developmental biologists are interested in the mechanisms that confer canalization; evolutionary biologists, however, are sought to understand its adaptive significance and how natural selection shapes canalized patterns (Gibson and Wagner 2000).

At the molecular level, factors such as hsp90 (Rutherford and Lindquist 1998), microRNAs (Posadas and Carthew 2014), and methylation (Wilkins 2005; Hallgrimsson et al. 2019) play crucial roles in buffering phenotypic variation. In our previous work, we systematically investigated the expression pattern of protein-coding genes and found that canalized protein-coding genes regulate caste differentiation in ants (Qiu et al. 2022). Given that proteins are the direct executors of phenotype, this canalized expression pattern is perhaps expected. Extending this perspective to the regulatory layer, our current findings reveal that lncRNAs exhibit comparable expression patterns and functional significance to protein-coding genes. This parallel suggests that canalization is not restricted to structural components but also extends to noncoding regulators that orchestrate gene expression. Such integration implies that lncRNAs participate in, and reinforce, the canalized genetic networks (GRNs) governing caste development, adding an additional layer of robustness to developmental control. From the evolutionary perspective, canalization reflects the reduction of phenotypic variability during development through stabilizing selection (Gibson and Wagner 2000). The canalized pattern of lncRNA implies that their regulation is also subject to such selection force. As lncRNAs have been integrated into the conserved caste determination pathways, it is reasonable to infer that the underlying GRNs governing caste differentiation are themselves canalized through development. This allows the system to achieve both robustness and conditional responsive to environmental or hormonal cues. JH treatment redirects lncRNA expression to induce alternative developmental trajectories, but the overall structure of the GRN remains canalized to prevent maladaptive phenotypes.

Therefore, the canalization of lncRNA expression provides an important molecular insight into how developmental robustness coexists with phenotypic plasticity in ants. It highlights a multilayered regulatory organization has evolved under stabilizing selection to maintain reliable caste differentiation, while retaining the flexibility necessary for environmental adaptation. The coexistence of robustness and plasticity may constitute a fundamental principle through which social insect development maintains evolutionary stability without losing developmental flexibility.

## Materials and Methods

### Sample collection

*M.pharaonis* used for tissue RNA-seq and functional experiment was derived from one colony collected from Xishuangbanna, Yunnan province, China and reared in the laboratory of Kunming institute of zoology. Colonies were maintained under a constant temperature of 27 ℃ and 65% humidity and kept in plastic boxes coated with fluon.

For tissue-specific RNA-seq samples, the newly-eclosed gynes within three days were collected and dissected in PBS. Five tissues, brain, mesosoma, fat body, midgut, and ovary, were used for RNA extraction with TRIzol Reagent (Invitrogen, 15596018). The young workers (with light color) were treated with the same method. Four worker tissues, brain, mesosoma, fat body, and midgut, were selected. Each sample only includes a single tissue from one individual. Then, the library construction and sequencing were performed by BGI, Shenzhen, China.

For JH treatment sampling, early 3^rd^ instar larvae were feed with Methoprene (JH workers) or ethanol (control workers/reproductives) for 3 times on day 1, 3, and 6 since the start of the experiment. Then, samples were collected for RNA-seq at five time points: day 2 (early 3^rd^ larvae), 5 (mid 3^rd^ larvae), 10 (late 3^rd^ larvae), prepupae, and young pupae. As the 3^rd^ larval period of reproductive is less than 10 days. Reproductive samples on day 6 were collected and their RNA-seq data were equivalent to that of worker on day 5 and 10. The detail experimental design was described in our previous study (Li et al. 2024).

### RNA extraction and RT-qPCR

Ant total RNA was extracted using TRIzol Reagent (Invitrogen, 15596018) following the manufacturer’s instructions. cDNA was generated from 1 μg total RNA using the PrimeScript RT reagent Kit with gDNA Eraser (Takara, RR047). After diluted 5-folds with distilled water, the cDNA was used as template of RT-qPCR. Then, RT-qPCR was conducted on Roche LightCycler96 with TB Green Premix Ex Taq II (Takara, RR820) using the settings as follows: a preincubation step at 95℃ for 5 min followed by 40 cycles of 3 step amplification for 20 s at 95℃, 20 s at 55℃, and 20 s at 72℃. ΔΔCt method was used to calculate the expression level of target gene with *EF1A* (*LOC105833278*) as reference gene. For the RT-qPCR conducted in tissue samples, we dissect brain, mesosoma, fat body, midgut, and ovary from gynes within 3 days. 20 individuals were pooled to extract RNA. In RNAi experiment for *ASfln*, the sex of larvae and prepupae was indistinguishable. *Male1* gene (*LOC105835607*), which show significantly higher expression in male, was used to identify the sex of the samples. The male individuals were removed when we calculate the RNAi efficiency. The primer sequences for RT-qPCR are listed in Dataset S5.

### dsRNA preparation and RNA interference

The fragments and primers for dsRNA were designed on the E-RNAi (www.dkfz.de/signaling/e-rnai3/) website. Then, dsRNA was synthesized using the MEGAscript RNAi kit (Invitrogen, AM1626) as described by the manufacturer. dsRNAs were dissolved in nuclease-free water with the final concentration of 8 μg/μL. A 484 bp fragment of eGFP was used as control to synthesize dsRNA of the same concentration. For *ASfln*, dsRNA was injected twice in late 3^rd^ instar larval and early pupal stages, respectively. The late 3^rd^ instar reproductive larvae were picked from their original colonies and immobilized using double-sided tapes. dsRNA was injected into the ventral side of cephalothorax junction with a micropipette needle prepared by a micropipette puller (P-2000, Sutter). Then the larvae were put into new queenless colonies with at least twice number of workers relative to the number of injected larvae. The 2^nd^ injection was conducted in early pupal stage with white body and compound eyes. dsRNA was injected into the terminal end of mesosoma. To improve the penetration efficiency to flight muscle, SPc was added to dsRNA (8 μg/μL of dsRNA: 1 μg/μL of SPc = 3:1; v/v) (Yan et al. 2021; Ma et al. 2022). Knockdown of *lncov* was performed in adult gynes. dsRNA was injected into the lateral tergum at the 5^th^, 7^th^ and 9^th^ day after eclosion. 8 μg/μL of dsRNA was used for both experimental and control groups in adult RNAi. The primer sequences for dsRNA synthesis are listed in Dataset S5.

### Ovary dissection and morphological measurements

To evaluate the efficacy of RNAi for *lncov* and *ASfln* knockdown, we conducted morphological measurements of focal traits. For *lncov*, ovaries of gynes were dissected in PBSTw (DEPC-treated PBS, 0.1% Tween20) under microscope, then the ovaries were transferred on an adhesive slide. Images were captured with stereomicroscope (SMZ18, Nikon) and the oocyte surface was measured by NIS-Elements. For measurements on adult gynes in the RNAi experiment of *ASfln*, all individuals were put on ice and anesthetized. Then they were properly positioned under the stereomicroscope, ensuring their bodies are vertical to eliminate errors caused by angular distortion. The steps of image capture and measurement were the same as described above.

### Hybridization Chain Reaction (HCR)

To investigate the spatial expression pattern of *lncov*, we conducted *in situ* hybridization to examine its expression in the ovary. Ovaries were dissected in PBSTw (1×PBS, 0.1% Tween20) and fixed in 4% PFA for 20 min. Then, the fixed ovaries were permeabilized in Detergent solution (1% SDS, 0.5% Tween20, 50mM pH7.5 Tris-HCl, 1mM pH8.0 EDTA, 150mM NaCl) for 20 min. Following three sequential washes with PBSTw (10 min, 5 min, and 5 min), the tissues were pre-incubated in 200 μL of probe hybridization buffer for 30 min at 37℃. The remaining procedures were performed in accordance with the standard HCR protocol for *Drosophila* embryos, as detailed on the MI website (https://www.molecularinstruments.com/). Image capture was performed using Nikon A1MP+ confocal microscopes. The probe sequences for HCR are listed in Dataset S5.

### Annotation of lncRNAs from transcriptomic data

We used the transcriptomic dataset that cover the whole embryonic stages to annotate lncRNA. RNAseq data were processed with SOAPnuke (https://github.com/BGI-flexlab/SOAPnuke) to filter low quality reads. High-quality PE reads (Q30>80) were mapped into reference genomes using Star (2.7.7) (Dobin et al. 2013) to obtain the genomic coordinates of the spliced transcripts in genomes. The genome assemblies we used were same as the previous study (Qiu et al. 2022): *Monomorium pharaonis* (NCBI accession GCF_013373865.1), *Acromyrmex echinatior* (In-house assembly provided by Globa Ant Genomics Alliance, In-house annotation with GeMoMa (version 1.7.1) (Keilwagen et al. 2016). Then we compared the coordinates information with Stringtie (v2.1.4) (Kovaka et al. 2019) and assembled the aligned reads into isoforms. To obtain the comprehensive transcript set, we combined transcripts across all developmental stages and the annotated lncRNA transcripts from NCBI. Screening by transcript abundance and length using the parameters (-T 0.5 -m 200 -c 0), we determined the final transcript datasets. We also removed the transcripts overlapping with protein-coding genes in the same strand with mRNA by gffcompare (v0.12.6) (Pertea and Pertea 2020).

To obtain a high-quality lncRNA dataset from developmental transcriptomes, we filtered the transcripts with coding potential in three ways: CPAT (v3.0.2) (Wang et al. 2013), CPC (v1.0.1) (Kong et al. 2007), and similarity with known proteins. CPAT assesses coding potential with an alignment-free logistic regression model. We selected the coding probability (CP) cutoff < 0.364 for the prediction. CPC predicts coding potential using sequence features and support vector machine. We used the default parameter to filter the transcripts with coding potential. Finally, we used BLAST to compare the transcripts with known proteins in Uniprot database with the parameters: -p blastx -e 1e-05. Only genes successfully passed our all three filters were included in our lncRNA annotation.

### Constructing the developmental trajectory network

For each species, we constructed a developmental trajectory network based on the transcriptomic similarities across all samples. We used the lncRNA expression matrix to construct a weighted undirected network. In this network, nodes correspond to samples, and edges represent the transcriptomic similarity between linked samples. The edges’ weights were calculated using Spearman’s correlation coefficient, indicating the extent of transcriptome similarity. To enhance the network’s overall signal, edges with weak connections were eliminated. Considering that lncRNA expression similarity between samples is generally lower than that of protein-coding genes, we set a threshold of 0.45 for weak connections, rather than the commonly suggested threshold of > 0.8 for anisogenic samples based on correlation coefficients.

Next, we used this adjacency matrix (the weighted undirected network) to visualize the developmental trajectory using the igraph package (v2.0.3) in R. This algorithm incorporates edge weights, ensuring that nodes with robust connections, which reflect high transcriptome abundance similarities, are clustered together. Edge colors were calibrated to their weights for enhanced visualization, with the color gradient ranging from white (weight = 0.45) to black (weight = 1).

### Quantification of caste developmental potential

Developmental potential (*ΔΔ*) of the target individual was calculated from the respective transcriptomic distance between the target individual and the two castes at the subsequent developmental stage. This score reflects the extent to which the target individual is likely to develop into a gyne or a (small) worker. The same calculation method is described in a previous study to quantify developmental potential based on protein-coding gene expression (Qiu et al. 2022).

In short, the transcriptional differences between stage t and t+1 are first removed with Combat package (from sva (v.3.44.0)) in R to singly focus on caste differences. The developmental potential of each individual (i) was then calculated with the following formula in our previous study (Qiu et al. 2022):

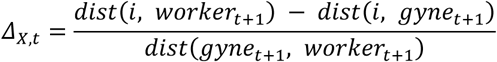

In this formula, dist (i, caste_t+1_) represents the Manhattan distance between individual i and the focal caste at the next stage. The difference in transcriptomic distances between castes was then normalized by dist (gyne_t+1_, worker_t+1_). A positive *Δ_X,t_* suggests that individual I is more likely to develop into a gyne, while a negative *Δ_X,t_* indicates a higher potential for developing into a worker.

### Detecting DEGs between castes

Raw reads from RNA-seq were filtered using trimmomatic (v0.39) (Bolger et al. 2014), and the clean reads were subsequently quantified by salmon (v1.10.2) (Patro et al. 2017), which generated both a read count matrix and a TPM matrix. Then the read count matrix was loaded into R using the tximport package (1.24.0) (Soneson et al. 2016), which integrated expression profiles from the transcript level to the gene level. Gene expression normalization and differential expression analysis were performed using DESeq2 (v1.36.0) (Love et al. 2014). Differentially expressed genes were identified based on the criteria of an absolute log2 (fold change) and Benjamini-Hochberg adjusted pvalue < 0.05. The raw reads from tissue-specific RNA-seq were treated following the same pipeline as described above.

### Caste tendency scores

This score is used to quantify the lncRNA profiles similarity of an individual to control gynes and workers. The formula to calculate caste tendency scores (CTS) is:

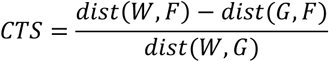

Dist means the Euclidean difference between different samples. F indicates a focal individual. W indicates mean transcriptome of same-staged gynes. G indicates mean transcriptome of same-staged gynes.

### Canalization-disruption score

The canalization-disruption score (*cd* score) reflects the effect of JH treatment on canalized expression pattern across developmental stages. Our previous study demonstrated protein-coding genes regulating caste differentiation undergo a canalized expression pattern across developmental stages, which means gradually increasing expression difference between castes and decreasing variation within caste (Qiu et al. 2022). Moreover, the expression pattern of some genes in JH workers redirecting from control workers to control gynes, mirroring the phenotypic changes of JH workers, indicating their contribution to caste differentiation process (Li et al. 2024). Here, we use the same logic to screen lncRNAs potentially relevant to caste development process. The lncRNAs with |*cd_l_*| > 1 and pvalue < 0.05 are defined as JH-responsive lncRNAs.

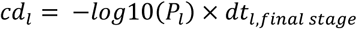

*P_l_* refers to the p value of the Pearson’s correlation coefficients to test if the *t_l_* increase over development. dt_l,final_ stage is the altered lncRNA expression difference between control workers and JH workers at the young pupal stage, such that:

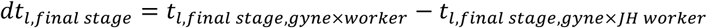

The *t* value of lncRNA *l* at developmental stage *s* between groups *a* and *b* can be formalized as follows:

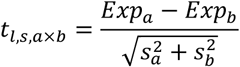

Where *Exp_a_* and *s_a_* represent the median and standard deviation of group *a*, respectively.

## Code availability

All code underlying these analyses and statistics in this study is available on Github (https://github.com/Ripcord-lin/AntslncRNA/tree/main).

## Acknowledgments

We thank Guangji Chen (Zhejiang University), Sheng Qian (Zhejiang University), Yaoxi He (Kunming institute of Zoology, Chinese Academy of Sciences), Yang Zhou (BGI), and members from Ehab Abouheif lab for helpful discussions. We also thank the Institutional Center for Shared Technologies and Facilities of Kunming Institute of Zoology (KIZ), Chinese Academy of Sciences (CAS) for technical support in Confocal Microscopy imaging. This work was supported by National Natural Science Foundation of China (Grant No. 32388102 to G.Z. and Grant No. 32370668 to W.L.), China Postdoctoral Science Foundation (Grant No. 2024M752769 to G.D.), Yunnan Provincial Science and Technology Department, Yunnan Fundamental Research Projects (Grant No. 202401BC070017 to W.L.), and New Cornerstone Science Foundation through the XPLORE PRIZE to G.Z..

## References

Abouheif E, Wray GA. 2002. Evolution of the Gene Network Underlying Wing Polyphenism in Ants. Science 297:249–252.

Adams RM, Larsen RS, Stylianidi N, Cheung D, Qiu B, Murray SK, Zhang G, Boomsma JJ. 2021. Hairs distinguish castes and sexes: identifying the early ontogenetic building blocks of a fungus-farming superorganism (Hymenoptera: Formicidae). Myrmecological News 31:201–216.

Bartel DP. 2009. MicroRNAs: target recognition and regulatory functions. Cell 136:215–233.

Bayala EX, Cisneros I, Massardo D, VanKuren NW, Kronforst MR. 2023. Divergent expression of *aristaless1* and *aristaless2* during embryonic appendage and pupal wing development in butterflies. BMC Biol 21:104.

Bekkevold D, Frydenberg J, Boomsma JJ. 1999. Multiple mating and facultative polygyny in the Panamanian leafcutter ant Acromyrmex echinatior. Behavioral Ecology and Sociobiology 46:103–109.

Bolger AM, Lohse M, Usadel B. 2014. Trimmomatic: a flexible trimmer for Illumina sequence data. Bioinformatics 30:2114–2120.

Bonasio R. 2012. Emerging topics in epigenetics: ants, brains, and noncoding RNAs. Annals of the New York Academy of Sciences 1260:14–23.

Bonasio R. 2014. The role of chromatin and epigenetics in the polyphenisms of ant castes. Briefings in Functional Genomics 13:235–245.

Bonasio R, Li Q, Lian J, Mutti NS, Jin L, Zhao H, Zhang P, Wen P, Xiang H, Ding Y, et al. 2012. Genome-wide and Caste-Specific DNA Methylomes of the Ants *Camponotus floridanus* and *Harpegnathos saltator*. Current Biology 22:1755–1764.

Bonasio R, Zhang G, Ye C, Mutti NS, Fang X, Qin N, Donahue G, Yang P, Li Q, Li C, et al. 2010. Genomic Comparison of the Ants *Camponotus floridanus* and *Harpegnathos saltator*. Science 329:1068–1071.

Boomsma JJ, Brady SG, Dunn RR, Gadau J, Heinze J, Keller L, Moreau CS, Sanders NJ, Schrader L, Schultz TR, et al. 2017. The Global Ant Genomics Alliance (GAGA). Myrmecological News 25:61–66.

Boomsma JJ, Gawne R. 2018. Superorganismality and caste differentiation as points of no return: how the major evolutionary transitions were lost in translation. Biological Reviews 93:28–54.

Campbell G, Tomlinson A. 1998. The roles of the homeobox genes *aristaless* and *Distal-less* in patterning the legs and wings of *Drosophila*. Development 125:4483–4493.

Chang M, Cheng H, Cai Z, Qian Y, Zhang K, Yang L, Ma N, Li D. 2022. miR-92a-1-p5 Modulated Expression of the *flightin* Gene Regulates Flight Muscle Formation and Wing Extension in the Pea Aphid, *Acyrthosiphon pisum* (Hemiptera: Aphidoidea). Mohamed A, editor. Journal of Insect Science 22:14.

Chen X, Shi W. 2020. Genome-wide characterization of coding and non-coding RNAs in the ovary of honeybee workers and queens. Apidologie 51:777–792.

Collins DH, Wirén A, Labédan M, Smith M, Prince DC, Mohorianu I, Dalmay T, Bourke AFG. 2021. Gene expression during larval caste determination and differentiation in intermediately eusocial bumblebees, and a comparative analysis with advanced eusocial honeybees. Molecular Ecology 30:718–735.

Debat V, Le Rouzic A. 2019. Canalization, a central concept in biology. Seminars in Cell & Developmental Biology 88:1–3.

Derrien T, Johnson R, Bussotti G, Tanzer A, Djebali S, Tilgner H, Guernec G, Martin D, Merkel A, Knowles DG, et al. 2012. The GENCODE v7 catalog of human long noncoding RNAs: Analysis of their gene structure, evolution, and expression. Genome Res. 22:1775–1789.

Dobin A, Davis CA, Schlesinger F, Drenkow J, Zaleski C, Jha S, Batut P, Chaisson M, Gingeras TR. 2013. STAR: ultrafast universal RNA-seq aligner. Bioinformatics 29:15–21.

Du H, Ge R, Zhang L, Zhang J, Chen K, Li C. 2023. Transcriptome-wide identification of development related genes and pathways in Tribolium castaneum. Genomics 115:110551.

Fandino RA, Brady NK, Chatterjee M, McDonald JMC, Livraghi L, Van Der Burg KRL, Mazo-Vargas A, Markenscoff-Papadimitriou E, Reed RD. 2024. The *ivory* lncRNA regulates seasonal color patterns in buckeye butterflies. Proc. Natl. Acad. Sci. U.S.A. 121:e2403426121.

Fatica A, Bozzoni I. 2014. Long non-coding RNAs: new players in cell differentiation and development. Nat Rev Genet 15:7–21.

Fernandes J, Acuña S, Aoki J, Floeter-Winter L, Muxel S. 2019. Long Non-Coding RNAs in the Regulation of Gene Expression: Physiology and Disease. ncRNA 5:17.

Ferreira PG, Patalano S, Chauhan R, Ffrench-Constant R, Gabaldón T, Guigó R, Sumner S. 2013. Transcriptome analyses of primitively eusocial wasps reveal novel insights into the evolution of sociality and the origin of alternative phenotypes. Genome Biol 14:R20.

Gao Q, Xiong Z, Larsen RS, Zhou L, Zhao J, Ding G, Zhao R, Liu C, Ran H, Zhang G. 2020. High-quality chromosome-level genome assembly and full-length transcriptome analysis of the pharaoh ant *Monomorium pharaonis*. GigaScience 9:giaa143.

Gibson G, Wagner G. 2000. Canalization in evolutionary genetics: a stabilizing theory? Bioessays 22:372–380.

Gospocic J, Glastad KM, Sheng L, Shields EJ, Berger SL, Bonasio R. 2021. Kr-h1 maintains distinct caste-specific neurotranscriptomes in response to socially regulated hormones. Cell 184:5807–5823.e14.

Hallgrimsson B, Green RM, Katz DC, Fish JL, Bernier FP, Roseman CC, Young NM, Cheverud JM, Marcucio RS. 2019. The developmental-genetics of canalization. Seminars in Cell & Developmental Biology 88:67–79.

Hölldobler B, Wilson EO. 1990. The ants. Harvard University Press

Hu W, Alvarez-Dominguez JR, Lodish HF. 2012. Regulation of mammalian cell differentiation by long non-coding RNAs. EMBO Reports 13:971–983.

Huang B, Zhang R. 2014. Regulatory non-coding RNAs: revolutionizing the RNA world. Mol Biol Rep 41:3915–3923.

Jean-Guillaume CB, Kumar JP. 2022. Development of the ocellar visual system in *Drosophila melanogaster*. The FEBS Journal 289:7411–7427.

Kapusta A, Kronenberg Z, Lynch VJ, Zhuo X, Ramsay L, Bourque G, Yandell M, Feschotte C. 2013. Transposable Elements Are Major Contributors to the Origin, Diversification, and Regulation of Vertebrate Long Noncoding RNAs. Hoekstra HE, editor. PLoS Genet 9:e1003470.

Keilwagen J, Wenk M, Erickson JL, Schattat MH, Grau J, Hartung F. 2016. Using intron position conservation for homology-based gene prediction. Nucleic Acids Research 44:e89.

Khila A, Abouheif E. 2008. Reproductive constraint is a developmental mechanism that maintains social harmony in advanced ant societies. Proc. Natl. Acad. Sci. U.S.A. 105:17884–17889.

Khila A, Abouheif E. 2010. Evaluating the role of reproductive constraints in ant social evolution. Phil. Trans. R. Soc. B 365:617–630.

Kong L, Zhang Y, Ye Z-Q, Liu X-Q, Zhao S-Q, Wei L, Gao G. 2007. CPC: assess the protein-coding potential of transcripts using sequence features and support vector machine. Nucleic Acids Research 35:W345–W349.

Kopp F, Mendell JT. 2018. Functional Classification and Experimental Dissection of Long Noncoding RNAs. Cell 172:393–407.

Kovaka S, Zimin AV, Pertea GM, Razaghi R, Salzberg SL, Pertea M. 2019. Transcriptome assembly from long-read RNA-seq alignments with StringTie2. Genome Biol 20:278.

Lebedev E, Smutin D, Timkin P, Kotelnikov D, Taldaev A, Panushev N, Adonin L. 2024. The Eusocial Non-Code: Unveiling the Impact of Noncoding RNAs on Hymenoptera Eusocial Evolution. Non-coding RNA Research [Internet]. Available from: https://www.sciencedirect.com/science/article/pii/S2468054024001501

Li Q, Wang Z, Lian J, Schiøtt M, Jin L, Zhang P, Zhang Y, Nygaard S, Peng Z, Zhou Y, et al. 2014. Caste-specific RNA editomes in the leaf-cutting ant *Acromyrmex echinatior*. Nat Commun 5:4943.

Li R, Dai X, Zheng J, Larsen RS, Qi Y, Zhang X, Vizueta J, Boomsma JJ, Liu W, Zhang G. 2024. Juvenile hormone as key regulator for asymmetric caste differentiation in ants. Proceedings of the National Academy of Sciences 121:e2406999121.

Liao Q, Liu C, Yuan X, Kang S, Miao R, Xiao H, Zhao G, Luo H, Bu D, Zhao H, et al. 2011. Large-scale prediction of long non-coding RNA functions in a coding–non-coding gene co-expression network. Nucleic Acids Research 39:3864–3878.

Livraghi L, Hanly JJ, Evans E, Wright CJ, Loh LS, Mazo-Vargas A, Kamrava K, Carter A, Van Der Heijden ESM, Reed RD, et al. 2024. A long noncoding RNA at the *cortex* locus controls adaptive coloration in butterflies. Proc. Natl. Acad. Sci. U.S.A. 121:e2403326121.

Love MI, Huber W, Anders S. 2014. Moderated estimation of fold change and dispersion for RNA-seq data with DESeq2. Genome Biology 15:550.

Luo S, Lu JY, Liu L, Yin Y, Chen C, Han X, Wu B, Xu R, Liu W, Yan P, et al. 2016. Divergent lncRNAs Regulate Gene Expression and Lineage Differentiation in Pluripotent Cells. Cell Stem Cell 18:637–652.

Ma Z, Zheng Y, Chao Z, Chen H, Zhang Y, Yin M, Shen J, Yan S. 2022. Visualization of the process of a nanocarrier-mediated gene delivery: stabilization, endocytosis and endosomal escape of genes for intracellular spreading. Journal of Nanobiotechnology 20:124.

Martin A, Reed RD. 2010. *wingless* and *aristaless2* Define a Developmental Ground Plan for Moth and Butterfly Wing Pattern Evolution. Molecular Biology and Evolution 27:2864–2878.

Mattick JS, Amaral PP, Carninci P, Carpenter S, Chang HY, Chen L-L, Chen R, Dean C, Dinger ME, Fitzgerald KA, et al. 2023. Long non-coding RNAs: definitions, functions, challenges and recommendations. Nat Rev Mol Cell Biol 24:430–447.

Okwaro LA, Korb J. 2023. Epigenetic regulation and division of labor in social insects. Current Opinion in Insect Science 58:101051.

Oldroyd BP, Yagound B. 2021. The role of epigenetics, particularly DNA methylation, in the evolution of caste in insect societies. Phil. Trans. R. Soc. B 376:20200115.

Orr SE, Goodisman MA. 2023. Social insect transcriptomics and the molecular basis of caste diversity. Current Opinion in Insect Science 57:101040.

Pan Q, Darras H, Keller L. 2024. LncRNA gene *ANTSR* coordinates complementary sex determination in the Argentine ant. Sci. Adv. 10:eadp1532.

Patro R, Duggal G, Love MI, Irizarry RA, Kingsford C. 2017. Salmon provides fast and bias-aware quantification of transcript expression. Nat Methods 14:417–419.

Penick CA, Prager SS, Liebig J. 2012. Juvenile hormone induces queen development in late-stage larvae of the ant *Harpegnathos saltator*. Journal of Insect Physiology 58:1643–1649.

Pertea G, Pertea M. 2020. GFF Utilities: GffRead and GffCompare. F1000Research 9:ISCB Comm J.

Pontieri L, Linksvayer TA. 2021. Monomorium. In: Starr CK, editor. Encyclopedia of Social Insects. Cham: Springer International Publishing. p. 599–604. Available from: http://link.springer.com/10.1007/978-3-030-28102-1_171

Ponting CP, Haerty W. 2022. Genome-Wide Analysis of Human Long Noncoding RNAs: A Provocative Review. Annual Review of Genomics and Human Genetics 23:null.

Ponting CP, Oliver PL, Reik W. 2009. Evolution and Functions of Long Noncoding RNAs. Cell 136:629–641.

Posadas DM, Carthew RW. 2014. MicroRNAs and their roles in developmental canalization. Current Opinion in Genetics & Development 27:1–6.

Qian SH, Chen L, Xiong Y-L, Chen Z-X. 2022. Evolution and function of developmentally dynamic pseudogenes in mammals. Genome Biol 23:235.

Qiu B, Dai X, Li P, Larsen RS, Li R, Price AL, Ding G, Texada MJ, Zhang X, Zuo D, et al. 2022. Canalized gene expression during development mediates caste differentiation in ants. Nat Ecol Evol 6:1753–1765.

Quinn JJ, Chang HY. 2016. Unique features of long non-coding RNA biogenesis and function. Nat Rev Genet 17:47–62.

Rajakumar A, Pontieri L, Li R, Larsen RS, Vásquez-Correa A, Frandsen JKL, Rafiqi AM, Zhang G, Abouheif E. 2024. From Egg to Adult: A Developmental Table of the Ant *Monomorium pharaonis*. J Exp Zool Pt B 342:557–585.

Rauluseviciute I, Riudavets-Puig R, Blanc-Mathieu R, Castro-Mondragon JA, Ferenc K, Kumar V, Lemma RB, Lucas J, Chèneby J, Baranasic D, et al. 2024. JASPAR 2024: 20th anniversary of the open-access database of transcription factor binding profiles. Nucleic Acids Research 52:D174–D182.

Rutherford SL, Lindquist S. 1998. Hsp90 as a capacitor for morphological evolution. Nature 396:336–342.

Sarropoulos I, Marin R, Cardoso-Moreira M, Kaessmann H. 2019. Developmental dynamics of lncRNAs across mammalian organs and species. Nature 571:510–514.

Seki R, Li C, Fang Q, Hayashi S, Egawa S, Hu J, Xu L, Pan H, Kondo M, Sato T, et al. 2017. Functional roles of Aves class-specific cis-regulatory elements on macroevolution of bird-specific features. Nat Commun 8:14229.

Shi Y-Y, Zheng H-J, Pan Q-Z, Wang Z-L, Zeng Z-J. 2015. Differentially expressed microRNAs between queen and worker larvae of the honey bee (*Apis mellifera*). Apidologie 46:35–45.

Shields EJ, Sheng L, Weiner AK, Garcia BA, Bonasio R. 2018. High-Quality Genome Assemblies Reveal Long Non-coding RNAs Expressed in Ant Brains. Cell Rep 23:3078–3090.

Sieber KR, Dorman T, Newell N, Yan H. 2021. (Epi)Genetic Mechanisms Underlying the Evolutionary Success of Eusocial Insects. Insects 12:498.

Simola DF, Ye C, Mutti NS, Dolezal K, Bonasio R, Liebig J, Reinberg D, Berger SL. 2013. A chromatin link to caste identity in the carpenter ant *Camponotus floridanus*. Genome Res. 23:486–496.

Soneson C, Love MI, Robinson MD. 2016. Differential analyses for RNA-seq: transcript-level estimates improve gene-level inferences. Available from: https://f1000research.com/articles/4-1521

Statello L, Guo C-J, Chen L-L, Huarte M. 2021. Gene regulation by long non-coding RNAs and its biological functions. Nat Rev Mol Cell Biol 22:96–118.

Szathmáry E, Smith JM. 1995. The major evolutionary transitions. Nature 374:227–232.

Ulitsky I, Shkumatava A, Jan CH, Sive H, Bartel DP. 2011. Conserved Function of lincRNAs in Vertebrate Embryonic Development despite Rapid Sequence Evolution. Cell 147:1537–1550.

Vigoreaux JO, Saide JD, Valgeirsdottir K, Pardue ML. 1993. Flightin, a novel myofibrillar protein of *Drosophila* stretch-activated muscles. The Journal of cell biology 121:587–598.

Vizueta J, Xiong Z, Ding G, Larsen RS, Ran H, Gao Q, Stiller J, Dai W, Jiang W, Zhao J, et al. 2025. Adaptive radiation and social evolution of the ants. Cell 188:4828–4848.e25.

Waddington CH. 1942. Canalization of Development and the Inheritance of Acquired Characters. Nature 150:563–565.

Waddington CH. 1957. The Strategy of the Genes: A Discussion of Some Aspects of Theoretical Biology. Allen & Unwin

Wang C, Dickinson LK, Lehmann R. 1994. Genetics of *nanos* localization in *Drosophila*. Developmental Dynamics 199:103–115.

Wang L, Park HJ, Dasari S, Wang S, Kocher J-P, Li W. 2013. CPAT: Coding-Potential Assessment Tool using an alignment-free logistic regression model. Nucleic Acids Research 41:e74.

Wheeler WM. 1910. Ants: their structure, development and behavior. Columbia University Press

Wilkens M, Zimbelmann S, Roth F, Cartano J, Sayols S, Dejung M, Levin M, Butter F. 2025. Unraveling developmental gene regulation in holometabolous insects through comparative transcriptomics and proteomics. Commun Biol 8:980.

Wilkins J. 2005. Genomic imprinting and methylation: epigenetic canalization and conflict. Trends in Genetics 21:356–365.

Wojciechowski M, Lowe R, Maleszka J, Conn D, Maleszka R, Hurd PJ. 2018. Phenotypically distinct female castes in honey bees are defined by alternative chromatin states during larval development. Genome Res. 28:1532–1542.

Yan S, Ren B, Shen J. 2021. Nanoparticle-mediated double-stranded RNA delivery system: A promising approach for sustainable pest management. Insect Science 28:21–34.

Zhang X, Xu Y, Chen B, Kang L. 2020. Long noncoding RNA PAHAL modulates locust behavioural plasticity through the feedback regulation of dopamine biosynthesis. PLOS Genetics 16:e1008771.

