## Supplemental Figures for "lncRNAs contribute to caste differentiation as a regulatory layer in ants"

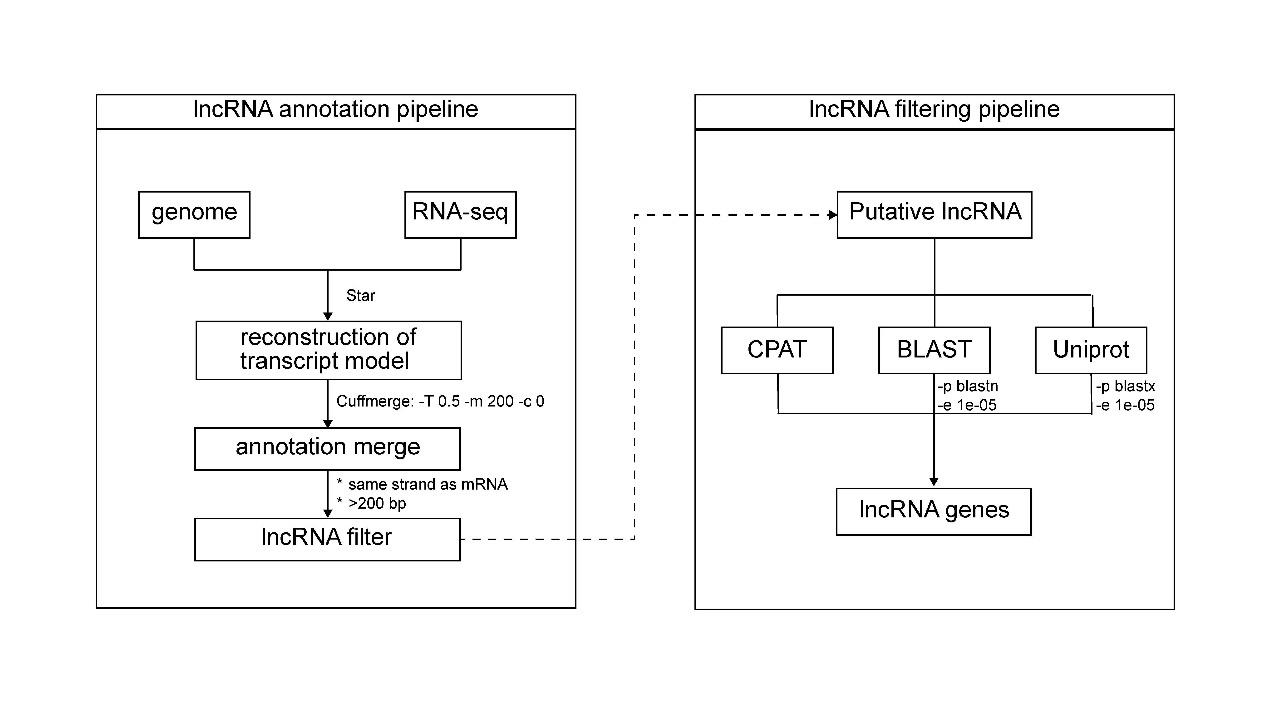

Fig. S1. Annotation and filtering of lncRNAs. Left panel illustrates the lncRNA annotation pipeline; right panel represents the lncRNA filtering pipeline. Softwares and thresholds utilized at each stage were also labeled.

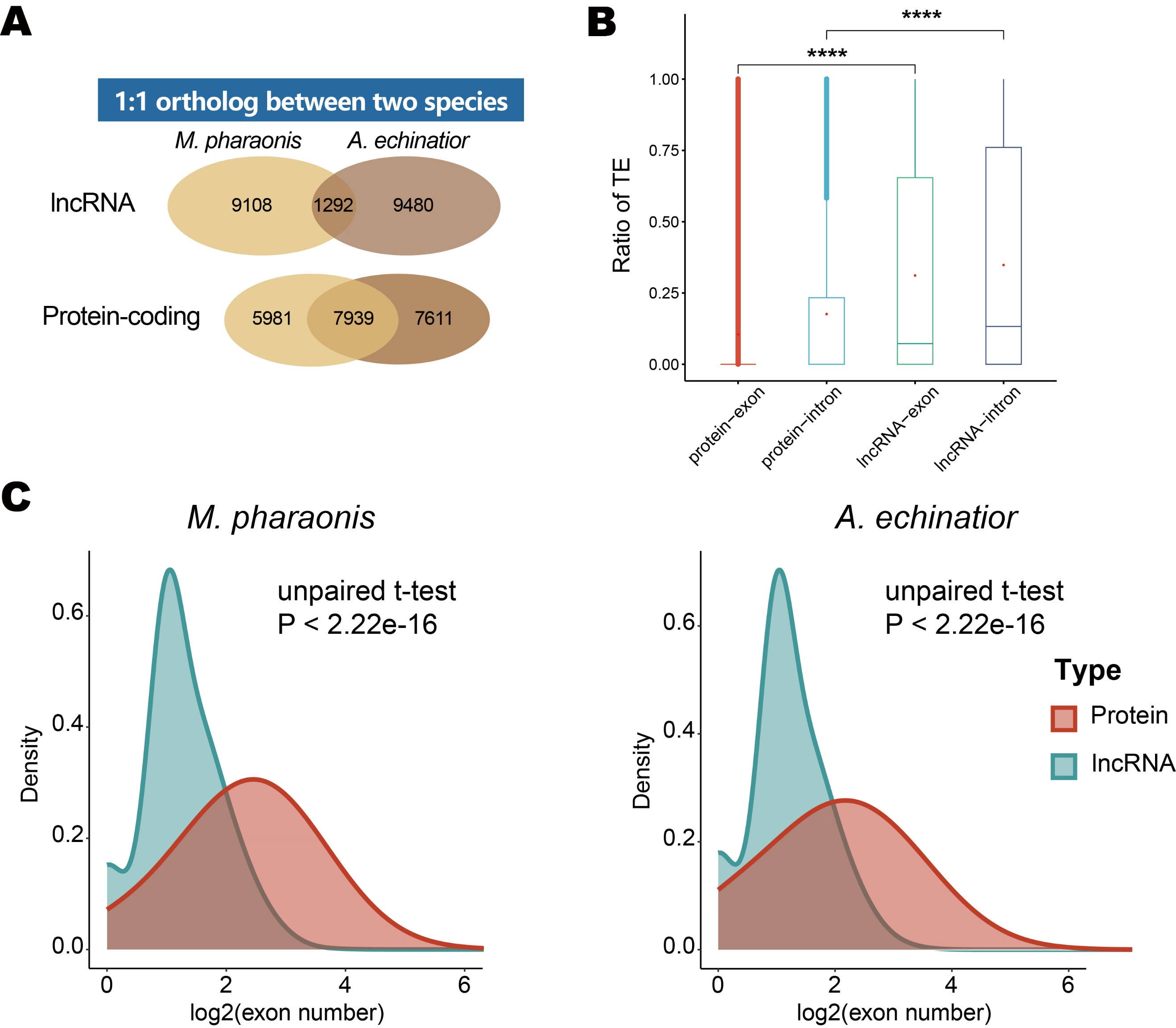

Fig. S2. Structural characters of lncRNAs in ants. (*A*) Numbers of lncRNAs and protein-coding genes with 1:1 ortholog in *M. pharaonis* and *A. echinatior*. Both lncRNAs and protein-coding genes were annotated with the same pipeline. Numbers in the overlapping parts between two species indicate the lncRNA numbers with ortholog. (*B*) Coverage of TE in different genomic elements (exon and intron) of protein-coding genes and lncRNAs. Each point represents the proportion of TE length in given regions (exon or intron), with comparison between the two gene types. The boxes represent the interquartile range (IQR), while error bars extend to the minimum and maximum values. pvalue < 2.2e-16 for both comparisons. (*C*) Density distribution of exon number for protein-coding genes and lncRNAs in *M. pharaonis* and *A. echinatior*. Comparisons show that lncRNAs have less exon number than protein-coding genes in both species. pvalue < 2.22e-16 for both comparisons.

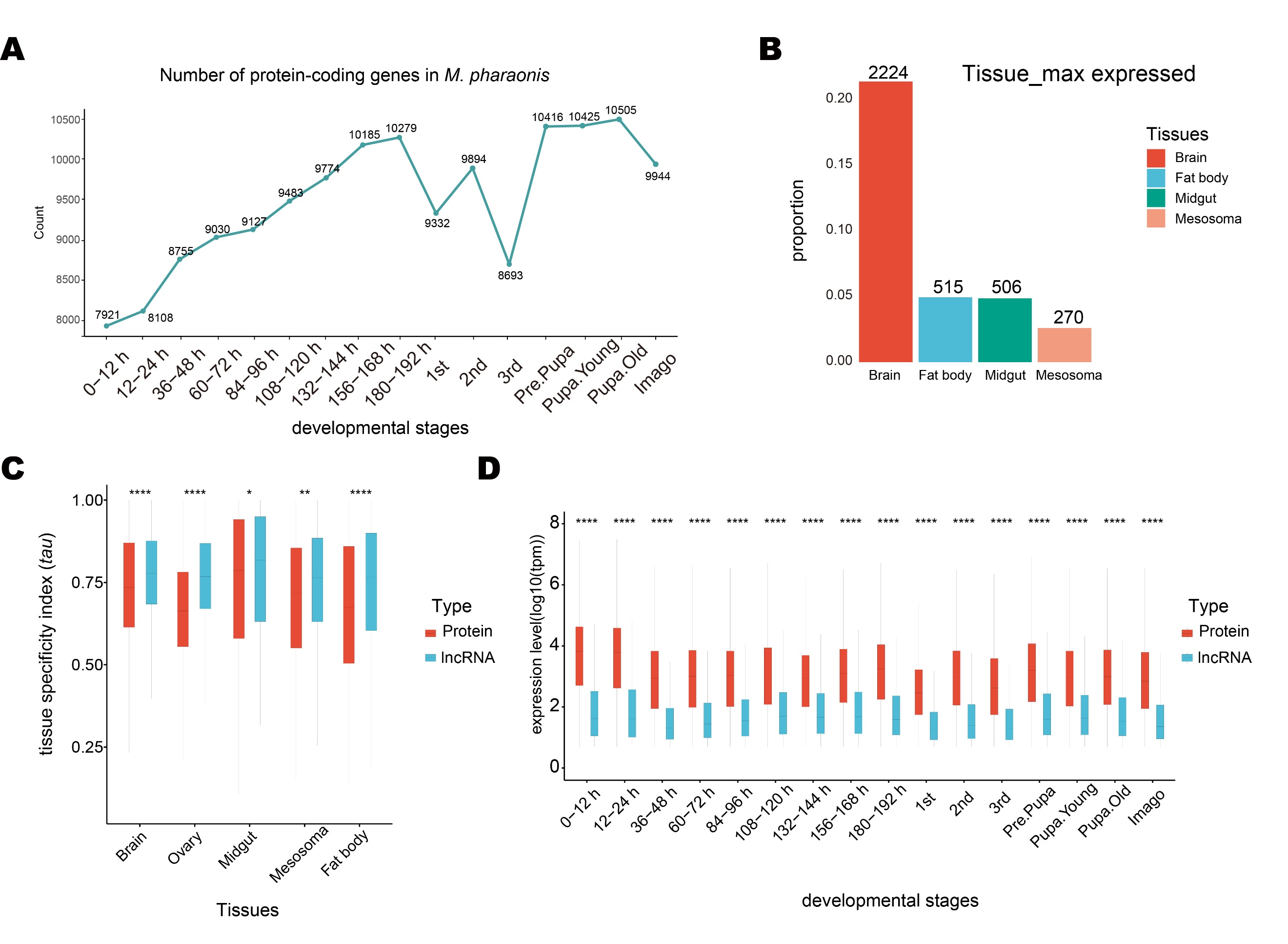

Fig. S3. Expression and distribution of lncRNAs in ants. (*A*) Number of expressed protein-coding genes (the average TPM > 1 in all samples) in each developmental stage in *M. pharaonis*. Notably, castes can be distinguished since the 2^nd^ instar larval stage, the count includes protein-coding genes from both castes (worker and gyne). (*B*) Number of lncRNAs with maximum expression in each tissue type. RNA-seq data from 4 tissues of adult young workers (with light body color) were used to calculate expression levels. Maximum expression refers to having the highest expression level in the samples of this tissue compared to other tissues. (*C*) Tissue-specificity comparison between protein-coding genes and lncRNAs across 5 tissues with RNA-seq data. Tissue-specificity is quantified by the tau value of each gene expressed in that tissue. The boxes represent the interquartile range (IQR), and error bars extend to the minimum and maximum values. pvalue < 2.2e-16 for brain, ovary, and fat body; pvalue = 0.0325 for midgut; pvalue = 0.0032 for mesosoma (t-test; * pvalue < 0.05; ** pvalue < 0.01; ***pvalue < 0.001; ****pvalue < 0.0001). (*D*) Comparison of expression levels between protein-coding genes and lncRNAs across each developmental stage. The boxes represent the interquartile range (IQR), and the error bars extend to the minimum and maximum values. pvalue < 2.2e-16 for all stages (t-test; * pvalue < 0.05; ** pvalue < 0.01; ***pvalue < 0.001; ****pvalue < 0.0001).

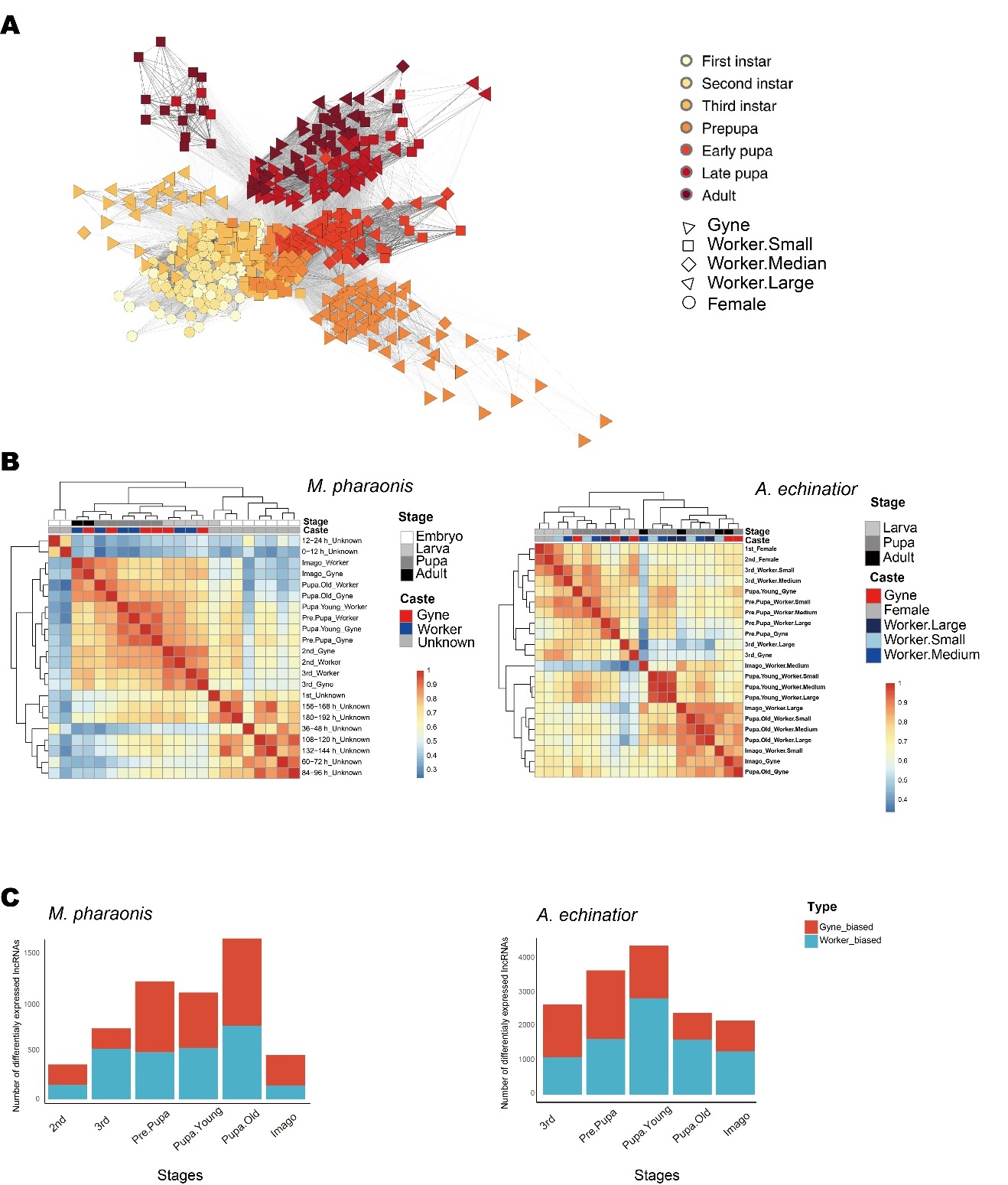

Fig. S4. Expression pattern of lncRNAs in ants. (*A*) Developmental trajectory of *A. echinatior* based on spearman correlation coefficients of lncRNA profiles between workers and gynes across morphologically distinguishable developmental stages. Each symbol represents an individual. 320 individuals were used in *A. echinatior*. (*B*) Between-stage lncRNA profiles similarity matrix across developmental stages in *M. pharaonis* (Left panel) and *A. echinatior* (Right panel) based on the mean values of within-group and between-group correlation coefficients. (*C*) Differentially expressed lncRNAs between gynes and (small) workers since morphologically distinguishable larval stage (2^nd^ and 3^rd^ for *M. pharaonis* and *A. echinatior*, respectively). Red represents gyne-biased lncRNAs and blue represents worker-biased lncRNAs.

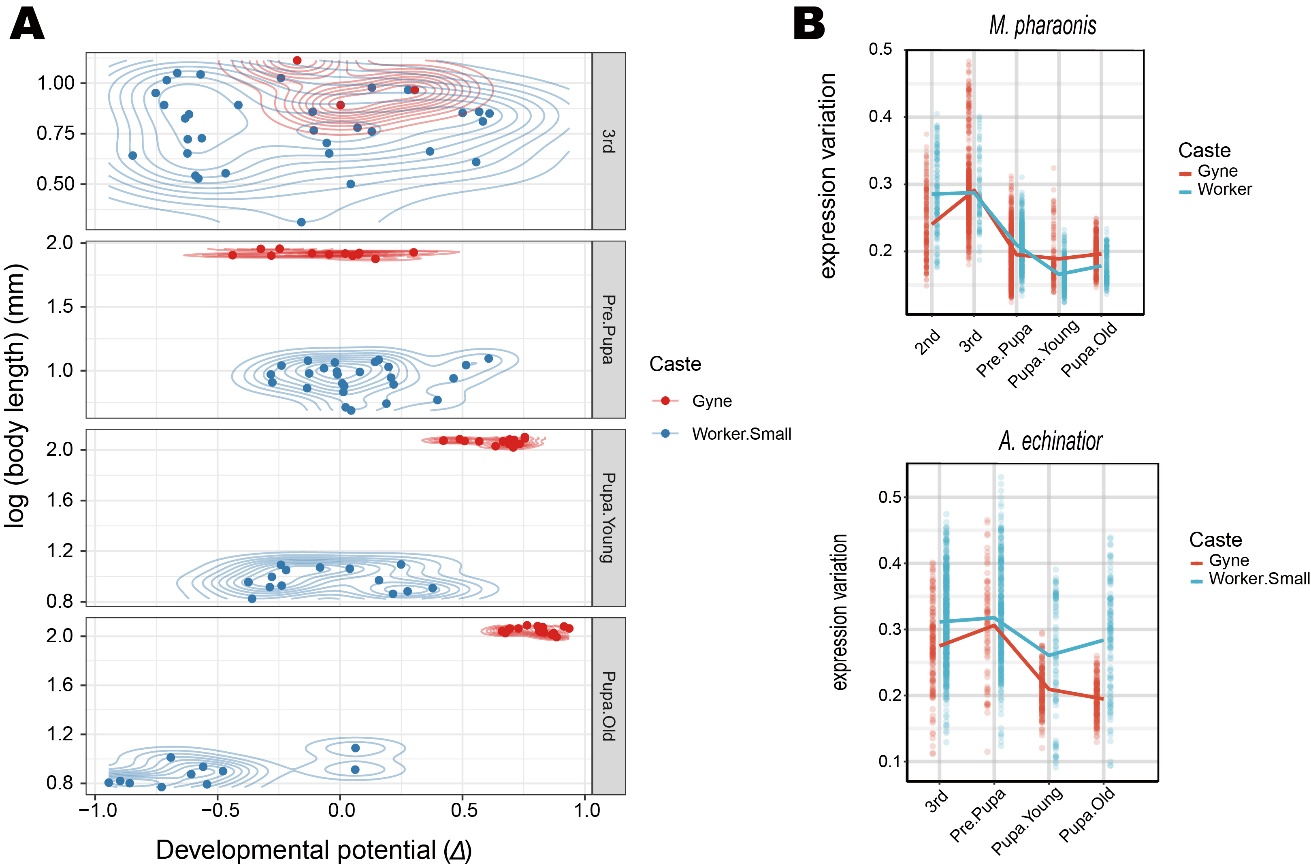

Fig. S5. Character of canalized lncRNA profile during caste differentiation in ants. (*A*) Developmental potential (*Δ*) score for individual gynes and workers in *A. echinatior*. The score was calculated since 3^rd^ instar larval stage as the workers of different subcastes were indistinguishable at 2^nd^ instar larval stage. (*B*) Expression variation between workers and gynes in each morphologically distinguishable developmental stage for *M. pharaonis* (Up panel) and *A. echinatior* (Down panel). The developmental stages used from the 1^st^ morphological distinguishable stage (2^nd^ and 3^rd^ instar larval stage for *M. pharaonis* and *A. echinatior*, respectively) to old pupal stage, when the morphological differentiation process between gynes and workers is largely completed.

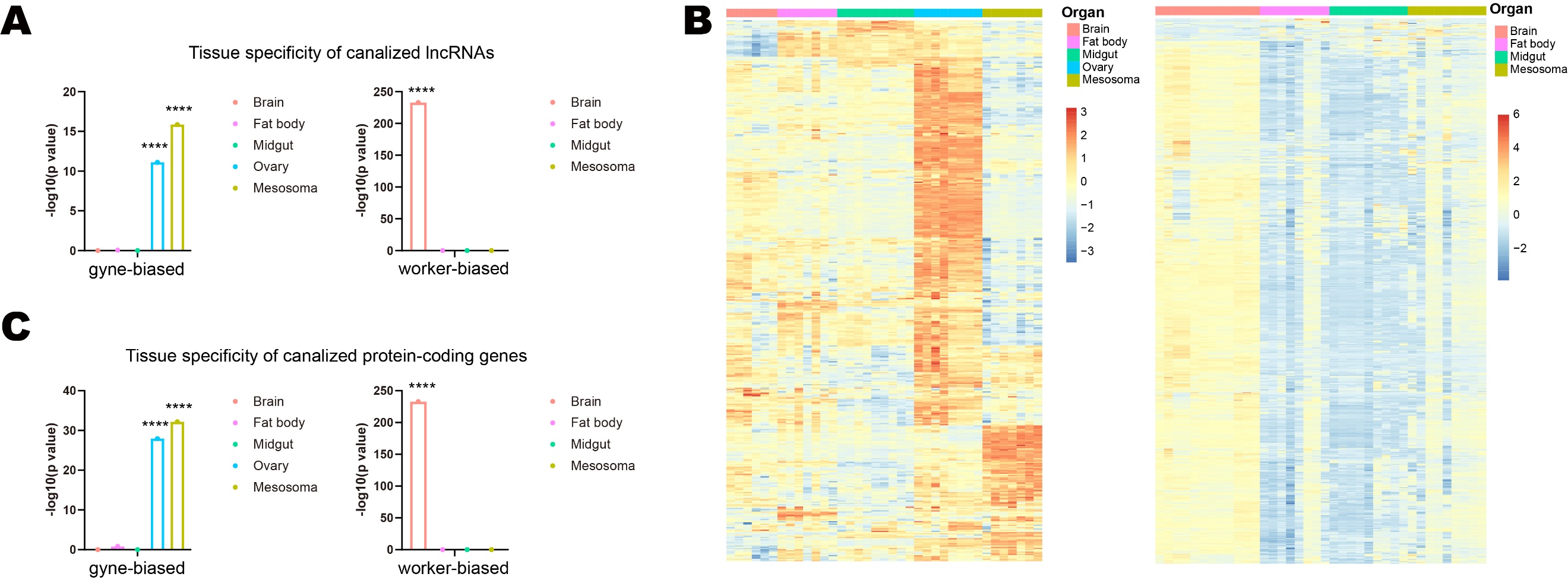

Fig. S6. Tissue distribution of protein-coding genes and lncRNAs in *M. pharaonis*. (*A*) The tissue specificity of canalized lncRNAs with gyne-biased (Left) and worker-biased (Right) expression patterns. The gyne-biased canalized lncRNAs enriched in ovary (pvalue = 7.76e-12, hypergeometric test; ****pvalue < 0.0001) and mesosoma (pvalue = 1.38e-16, hypergeometric test), while worker-biased canalized lncRNAs enriched in the brain (pvalue = 1.7e-233, hypergeometric test; ****pvalue < 0.0001). (*B*) Tissue-specific relative expression of canalized protein-coding genes with gyne-biased (Left) and worker-biased (Right) expression pattern. Heatmap shows expression value (TPM) for each protein-coding gene by row, with columns clustered by tissue and rows clustered by expression similarity. The expression values are normalized and shown as Z-scores. 5 adult gyne tissues, brain, fat body, midgut, ovary and mesosoma, were used for gyne-biased genes; while 4 adult worker tissues, brain, fat body, midgut and mesosoma, were used for worker-biased genes. Protein-coding gene lists were formed according to canalization score (n = 456 for gyne-biased, n = 748 for worker-biased, |C.score| ≥ 3, pvalue ≤ 0.05). (*C*) The tissue specificity of canalized protein-coding genes with gyne-biased (Left) and worker-biased (Right) expression patterns. The gyne-biased canalized genes enriched in ovary (pvalue = 1.14e-28, hypergeometric test; ****pvalue < 0.0001) and mesosoma (pvalue = 6.46e-33, hypergeometric test), while worker-biased canalized genes enriched in the brain (pvalue = 5.1e-159, hypergeometric test; ****pvalue < 0.0001).

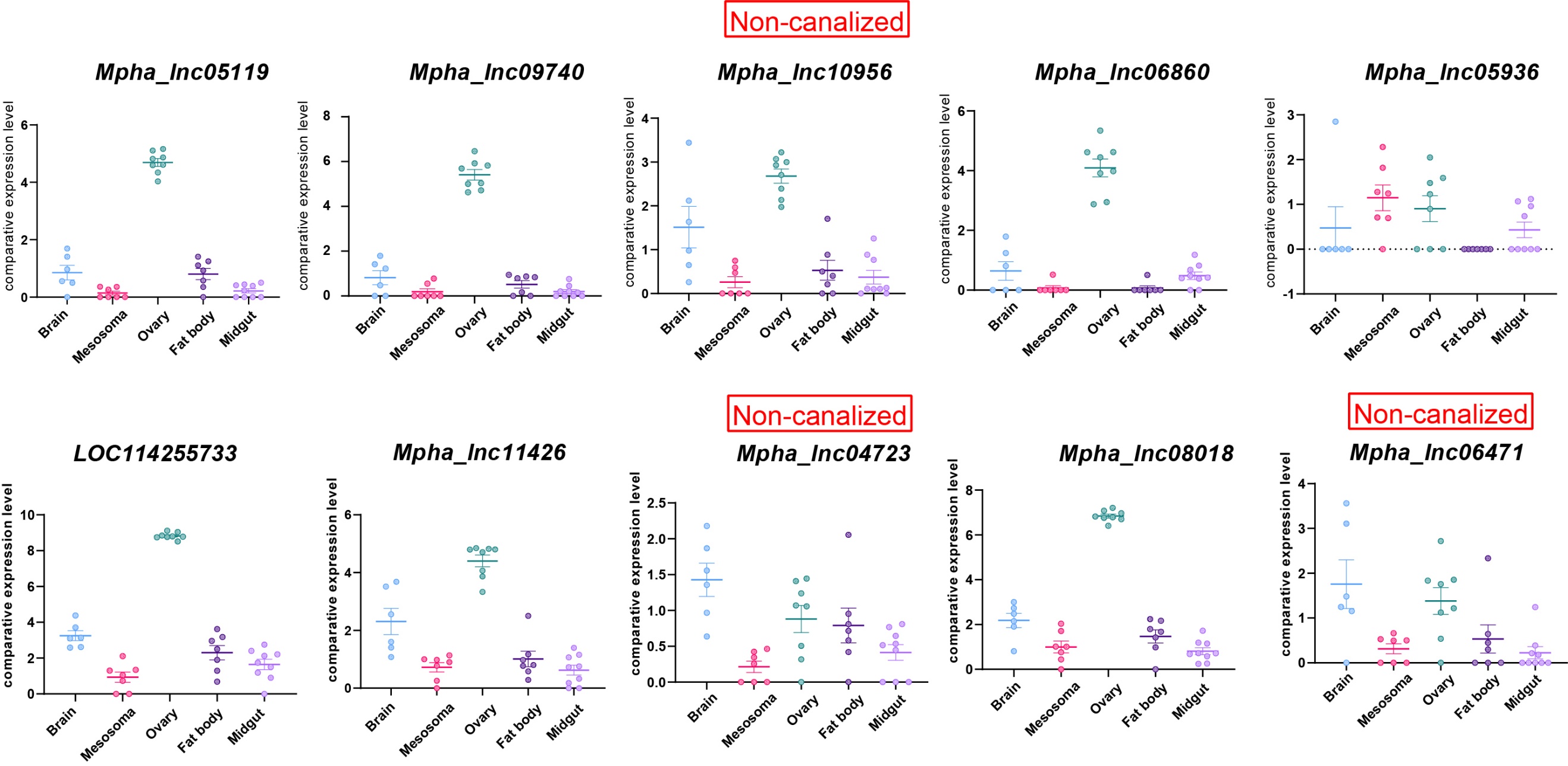

Fig. S7. Tissue-specificity of top 10 co-expressed lncRNAs with *Freja*, the top canalized protein-coding gene in *M. pharaonis*. The tissue distribution of these 10 lncRNAs, with expression shown as TPM. lncRNAs without canalization patterns are labeled as non-canalized (red). Canalized lncRNAs are predominantly enriched in ovary (except *Mpha_lnc05936*), similar to *Freja*; whereas non-canalized lncRNAs show no tissue specificity.

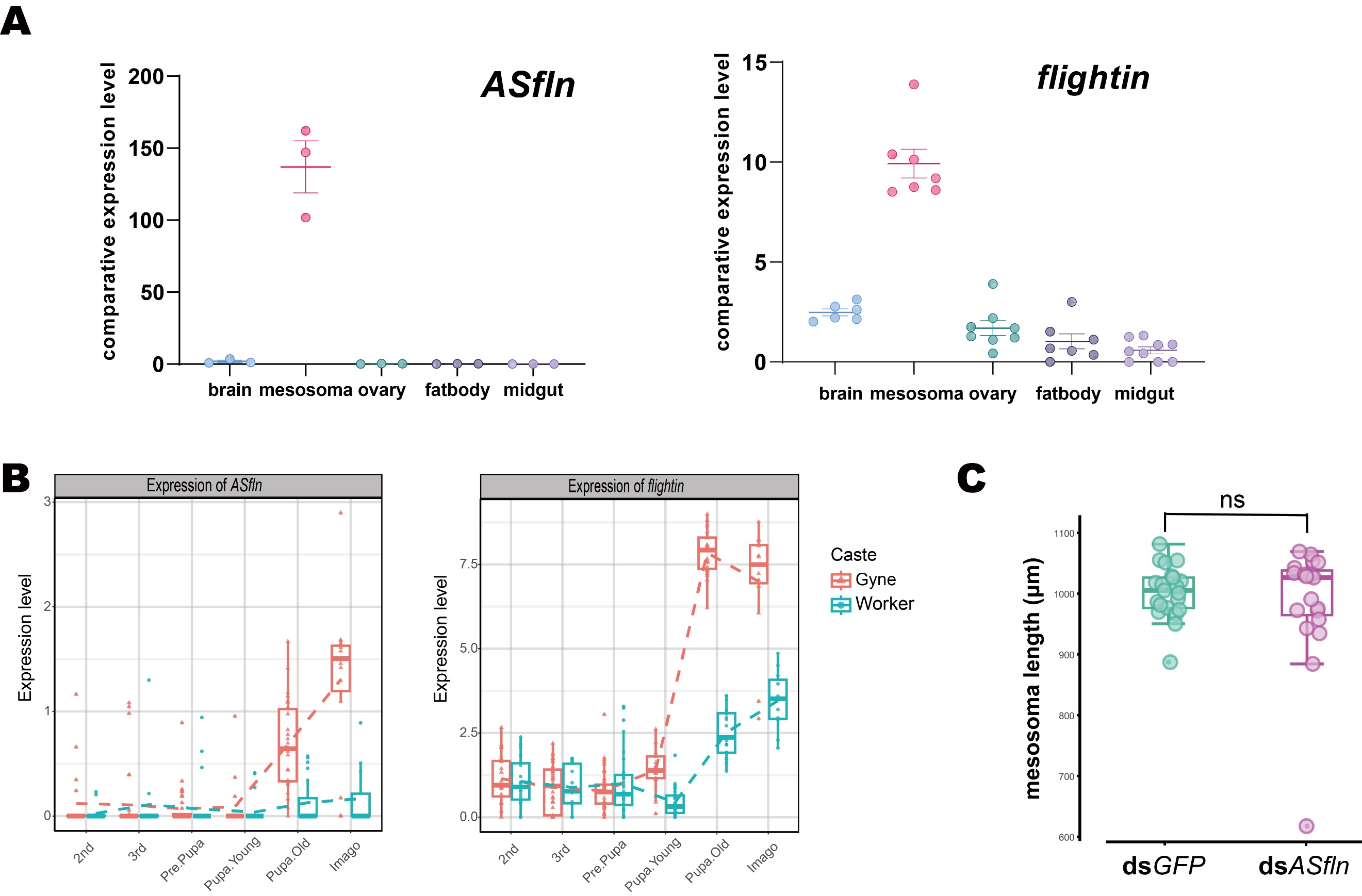

Fig. S8. Functional validation of *ASfln*. (*A*) Tissue distribution of *ASfln* and its antisense protein-coding gene *flightin* verified by RT-qPCR. Three biological replicates were used for RT-qPCR (mean ± SD). (*B*) The expression pattern of *ASfln* and *flightin* across developmental stages, which are drawn according to RNA-seq data. (*C*) Morphological measurements in RNAi experiment. The mesosoma length show no significant difference for two groups. pvalue = 0.4496 (two-tailed t-test, n = 21 for ds*GFP* and n = 19 for ds*ASfln*).

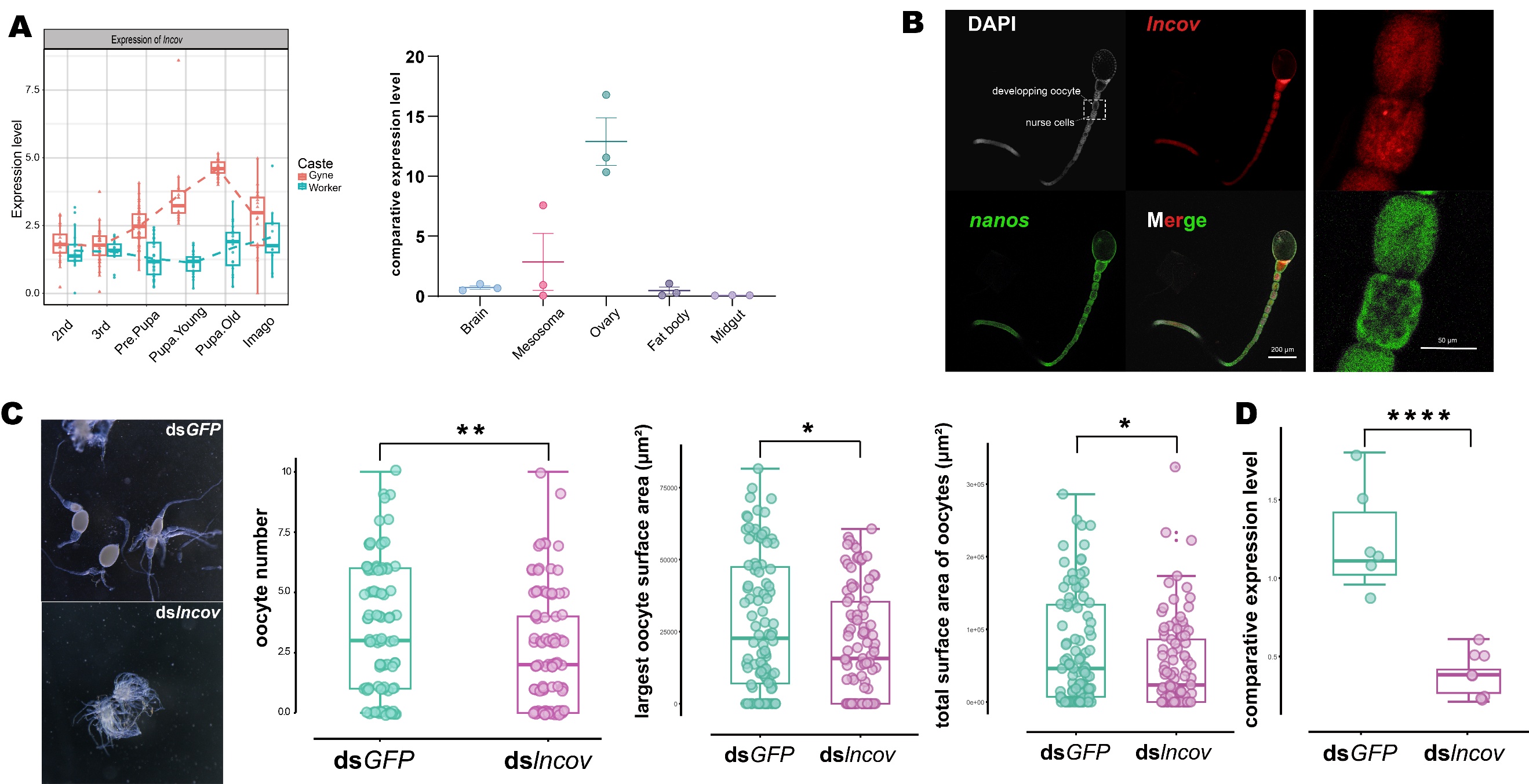

Fig. S9. Functional validation of *lncov*. (*A*) The expression pattern across developmental stages and tissue distribution of *lncov*. The expression pattern was drawn according to RNA-seq data. The tissue distribution was verified by RT-qPCR. Three biological replicates were used for RT-qPCR (mean ± SD). (*B*) HCR stain showing the localization of *lncov* in ovary. Left panel displays an intact ovariole and the area of the dotted box (including developing oocyte and nurse cells) enlarged in the right panel. Scale bar: 200 μm (left) and 50 μm (right). (*C*) RNAi phenotype. Left photos are images of ovaries from the two RNAi experimental groups. The ellipses with tail represent mature oocytes. Boxplot on the right shows the statistics of oocyte numbers (two-tailed t-test, pvalue = 0.0069), largest oocyte surface area (two-tailed t-test, pvalue = 0.0147), and total surface area of oocytes (two-tailed t-test, pvalue = 0.0292) between two groups. The boxes represent the interquartile range (IQR), while the error bars extend to the minimum and maximum values. (n = 94 for ds*GFP* and n = 99 for ds*lncov*). (*D*) Comparative expression level of *lncov* in the two RNAi experimental groups. (two-tailed t-test, pvalue = 0.0008 (n = 6 for ds*GFP* and n = 6 for ds*lncov*)). RT-qPCR was conducted after 48 h of injection.

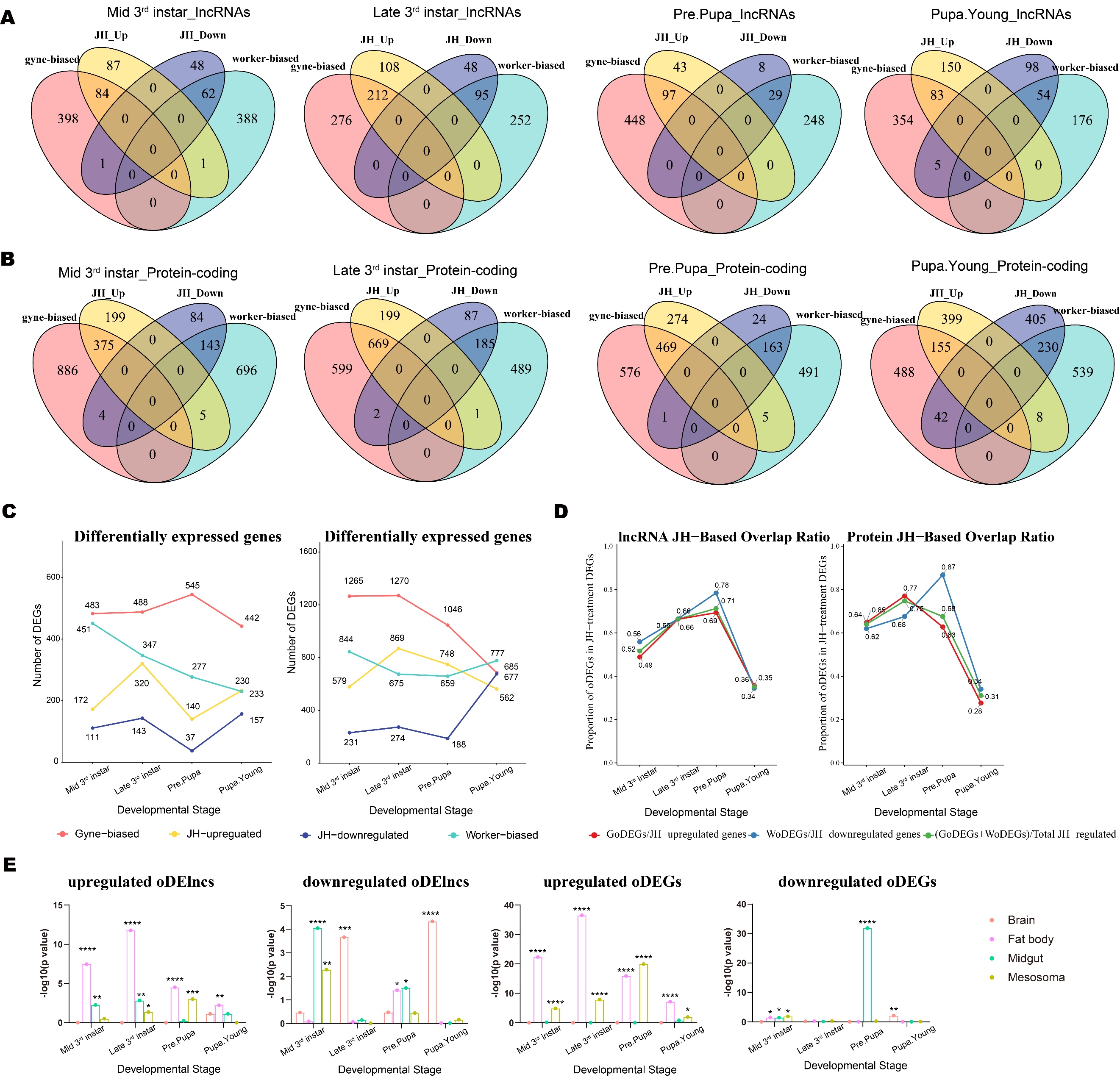

Fig. S10. Caste- and JH-responsive DEGs and DElncs across the developmental stages. (*A*) Venn diagrams showing numbers of caste-biased differentially expressed lncRNAs (DElncs) and JH-responsive lncRNAs in each stage. Red and blue represent the gyne-/worker-biased DElncs, respectively. Yellow and purple represent the JH-upregulated/downregulated DElncs, respectively. (*B*) Venn diagrams showing numbers of caste-biased differentially expressed protein-coding genes (DEGs) and JH-responsive protein-coding genes in each stage. Red and blue represent the gyne-/worker-biased DElncs, respectively. Yellow and purple represent the JH-upregulated/downregulated DElncs, respectively. (*C*) Numbers of caste-biased (gyne/worker) differentially expressed and JH-responsive (up/down) lncRNAs (Left) and protein-coding genes (Right) in each stage. (*D*) Proportion of overlapping DElncs/DEGs (oDElncs/oDEGs) in JH responsive lncRNAs (Left) and protein-coding genes (Right). Red line represents the gyne-biased overlapping differentially expressed lncRNAs/protein-coding genes. (*E*) Tissue-specificity of oDElncs and oDEGs (hypergeometric test; * pvalue < 0.05; ** pvalue < 0.01; ***pvalue < 0.001; ****pvalue < 0.0001).

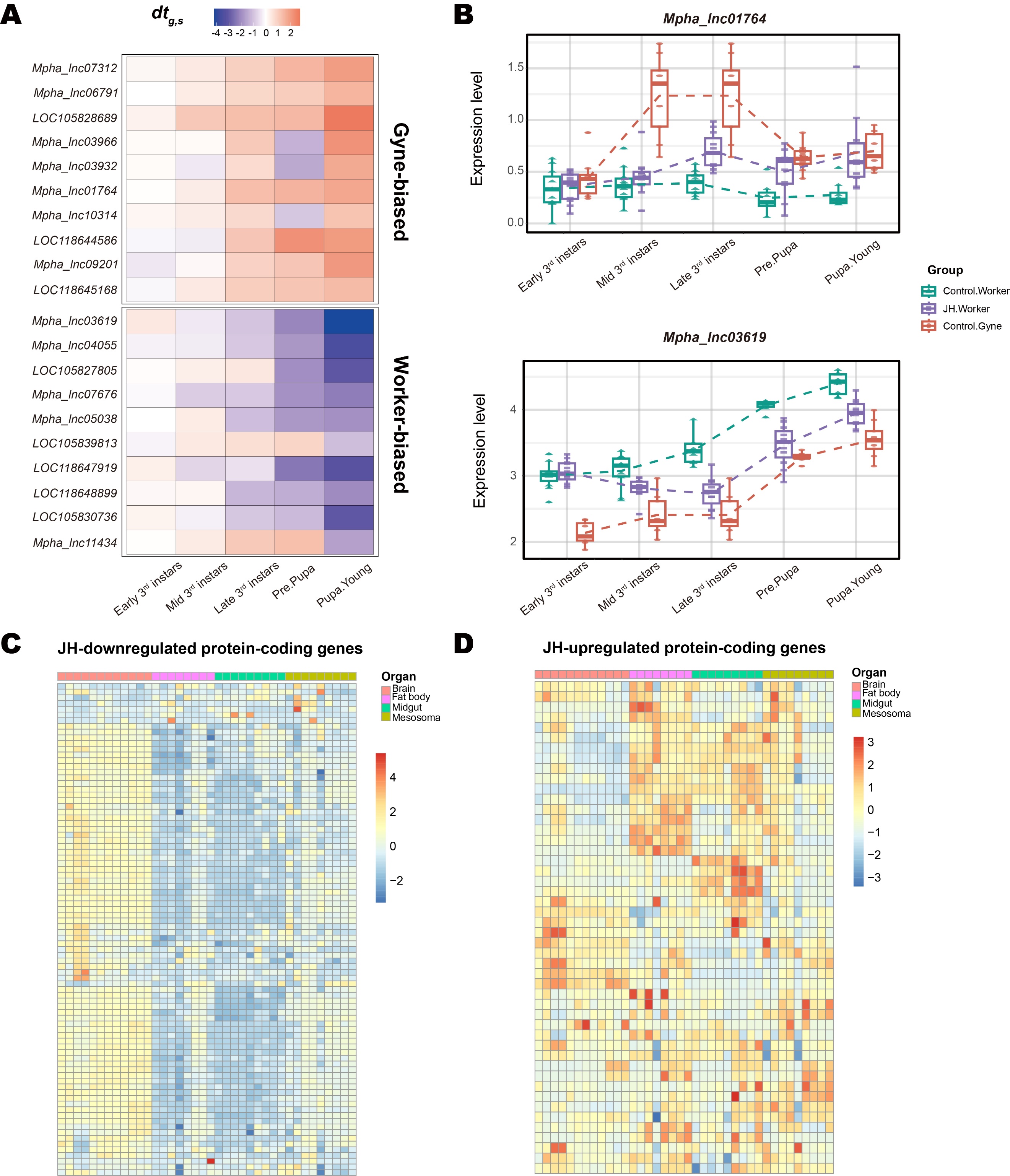

Fig. S11. The expression patterns of representative JH-responsive lncRNAs and tissue distribution of JH-responsive protein-coding genes. (*A*) The gene list is formed according to canalization-disruption (*cd*) scores. The heatmap colors represent *dt_g,s_* values, which reflecting the extent of JH treatment effects in each stage (Materials and Methods). (*B*) Two examples of gyne-biased and worker-biased lncRNAs, whose expression patterns were redirected to the other caste during development. (*C*) Tissue-specific relative expression of protein-coding genes down-regulated by JH in *M. pharaonis*. Heatmap shows expression value (TPM) for each lncRNA by row, with columns clustered by tissue and rows clustered by expression similarity. The expression values are normalized and shown as Z-scores, 4 worker tissues, brain, fat body, midgut and mesosoma, were used. (*D*) Tissue-specific relative expression of protein-coding genes up-regulated by JH in *M. pharaonis*. Heatmap shows expression value (TPM) for each lncRNA by row, with columns clustered by tissue and rows clustered by expression similarity. The expression values are normalized and shown as Z-scores, 4 worker tissues, brain, fat body, midgut and mesosoma, were used.

Dataset S1 (separate file). lncRNAs with canalized expression patterns in both ant species. This dataset includes 823 and 185 canalized lncRNAs in *M. pharaonis* and *A. echinatior*, respectively. In *M. pharaonis*, 62 lncRNAs are gyne-biased and 761 lncRNAs are worker-biased. In *A. echinatior*, 125 lncRNAs are gyne-biased and 60 lncRNAs are small worker-biased.

Dataset S2 (separate file). DElncs and DEGs between control castes and JH-regulated at 5 time points. In this dataset, each sheet represents the gene list of one time point for lncRNA or protein-coding genes. Column A and D are gene lists derived from comparisons between the control castes, whereas column G and J are gene lists from comparisons between control workers and JH workers. Genes highlight in red represent gyne-biased genes that overlap between the control caste comparison and the JH treatment comparison, while genes highlighted in green represent worker-biased genes overlapping between these two comparisons.

Dataset S3 (separate file). JH-responsive lncRNAs in *M. pharaonis*. This dataset represents the *cd* scores of all lncRNAs. JH-responsive lncRNAs screened with pvalue < 0.05 and |cd| ≥ 1 are highlighted in red. 168 lncRNAs are JH-upregulated, while 149 lncRNAs are JH-downregulated.

Dataset S4 (separate file). Motif enrichment for gyne-biased lncRNAs regulated by JH. In this dataset, the *motif enrichment* sheet lists all motifs enriched at a significance threshold of pvalue < 0.5, whereas the *potential function* sheet summarizes the detailed biological functions of the most significant transcription factors from uniprot and flybase databases.

Dataset S5 (separate file). Primer sequences used in this study. This dataset includes all primer sequences used for RT-qPCR and gene amplification, as well as the HCR probe sequences.
